# Beyond sequence similarity: cross-phyla protein annotation by structural prediction and alignment

**DOI:** 10.1101/2022.07.05.498892

**Authors:** Fabian Ruperti, Nikolaos Papadopoulos, Jacob Musser, Milot Mirdita, Martin Steinegger, Detlev Arendt

**Author notes:** These authors contributed equally to this work (Fabian Ruperti, Nikolaos Papadopoulos).

## Abstract

**Background:** Annotating protein function is a major goal in molecular biology, yet experimentally determined knowledge is often limited to a few model organisms. In non-model species, the sequence-based prediction of gene orthology can be used to infer function, however this approach loses predictive power with longer evolutionary distances. Here we propose a pipeline for the functional annotation of proteins using structural similarity, exploiting the fact that protein structures are directly linked to function and can be more conserved than protein sequences.

**Results:** We propose a pipeline of openly available tools for the functional annotation of proteins via structural similarity (MorF: **Mor**pholog**F**inder) and use it to annotate the complete proteome of a sponge. Sponges are highly relevant for inferring the early history of animals, yet their proteomes remain sparsely annotated. MorF accurately predicts the functions of proteins with known homology in *>*90% cases, and annotates an additional 50% of the proteome beyond standard sequence-based methods. Using this, we uncover new functions for sponge cell types, including extensive FGF, TGF and Ephrin signalling in sponge epithelia, and redox metabolism and control in myopeptidocytes. Notably, we also annotate genes specific to the enigmatic sponge mesocytes, proposing they function to digest cell walls.

**Conclusions:** Our work demonstrates that structural similarity is a powerful approach that complements and extends sequence similarity searches to identify homologous proteins over long evolutionary distances. We anticipate this to be a powerful approach that boosts discovery in numerous -omics datasets, especially for non-model organisms.

## Background

Knowledge of protein function is crucial for interpreting many types of high-throughput molecular datasets. Since protein functional studies are limited to a few model species, amino acid sequence similarity has been used to predict the function of protein homologs [1, 2].

However, homology detection over longer evolutionary distances remains challenging owing to the decay of protein sequence similarity that abolishes evidence of historical continuity. This presents a severe bottleneck for inferring protein function across a wide expanse of the tree of life, particularly in distant organisms where many proteins fall in the “twilight zone”, only sharing a sequence identity between 10 *−* 20% with proteins in characterised models [3, 4].

A way to venture more deeply into the twilight zone is to use structural similarity for homology detection, as structure is more conserved in evolution [5]. Until recently, this was not feasible since predicting protein structures from amino acid sequence required prior inference of sequence homology [6]. This has changed with the advent of AlphaFold [7], a deep learning AI system that can predict *de novo* protein structures with atomic resolution, together with novel approaches for identifying similar structures in large databases [8]. Protein structures can now be predicted by AlphaFold for entire proteomes, and then aligned to structures from model systems with characterised functions.

Sponges (*Porifera*) are animals that diverged early in the Metazoan tree relative to well annotated model organisms such as human and mouse. In this work, we predicted structures for the sparsely annotated proteome of the freshwater sponge *Spongilla lacustris* and aligned them against available structural databases to identify structurally similar proteins which we termed *“morpholog”* (from Greek *morphé* “form” and *lógos* “ratio”). We show that morphologs reflect homologous proteins in the vast majority of cases and often overlap in function even when homology is no longer detectable. We use morphologs to predict functions for unannotated sponge proteins by transferring functional annotations from model species. This complements sequence-based homology detection and subsequent function prediction. This expands the functional annotation coverage of the *Spongilla* proteome by 50%. Revisiting recent single-cell RNA-sequencing data [9] the novel annotations suggest additional aspects of sponge cell biology, such as extended cell signalling in pinacocytes, redox metabolism and control in myopeptidocytes, and polysaccharide digestion as a key function of the previously uncharacterised mesocytes.

### Results

#### A protein structure-based pipeline enriches functional annotation for *Spongilla lacustris*

We created a structure-based pipeline for functional annotation transfer, which we refer to as MorF (MorF: **Mor**pholog**F**inder). Instead of using amino acid sequence similarity to assign homology and predict function, we predict protein structures, align them against structural databases, and transfer the functional annotation of the best hits, such as their preferred name and description, to the queries (for an overview, see Suppl. Fig. S1; details in Methods). As a test case, we chose to annotate proteins in the freshwater demosponge *Spongilla lacustris*, an early-branching animal. With only about 20 cell types, organised into four cell families, *Spongilla* is a key model for understanding the origins of specialised animal cells [9].

We used the ColabFold [10] pipeline to predict three-dimensional structures for all 41,943 predicted *Spongilla* proteins, including isoforms (all structures and metadata deposited to ModelArchive [11], see Methods). Eleven (11) proteins were too long (*>*2, 900 amino acids) to be predicted by the available hardware, and were left unresolved. Confidence of predicted protein structures was assessed by calculating average pLDDT (predicted local distance difference) values. Average pLDDT values for *Spongilla* predicted protein structures were 4-6 percentage points lower than those of well-characterised animal models (Fig. 1A, [12]), likely reflecting the underrepresentation of sponges in the protein structure databases.

**Fig 1.**
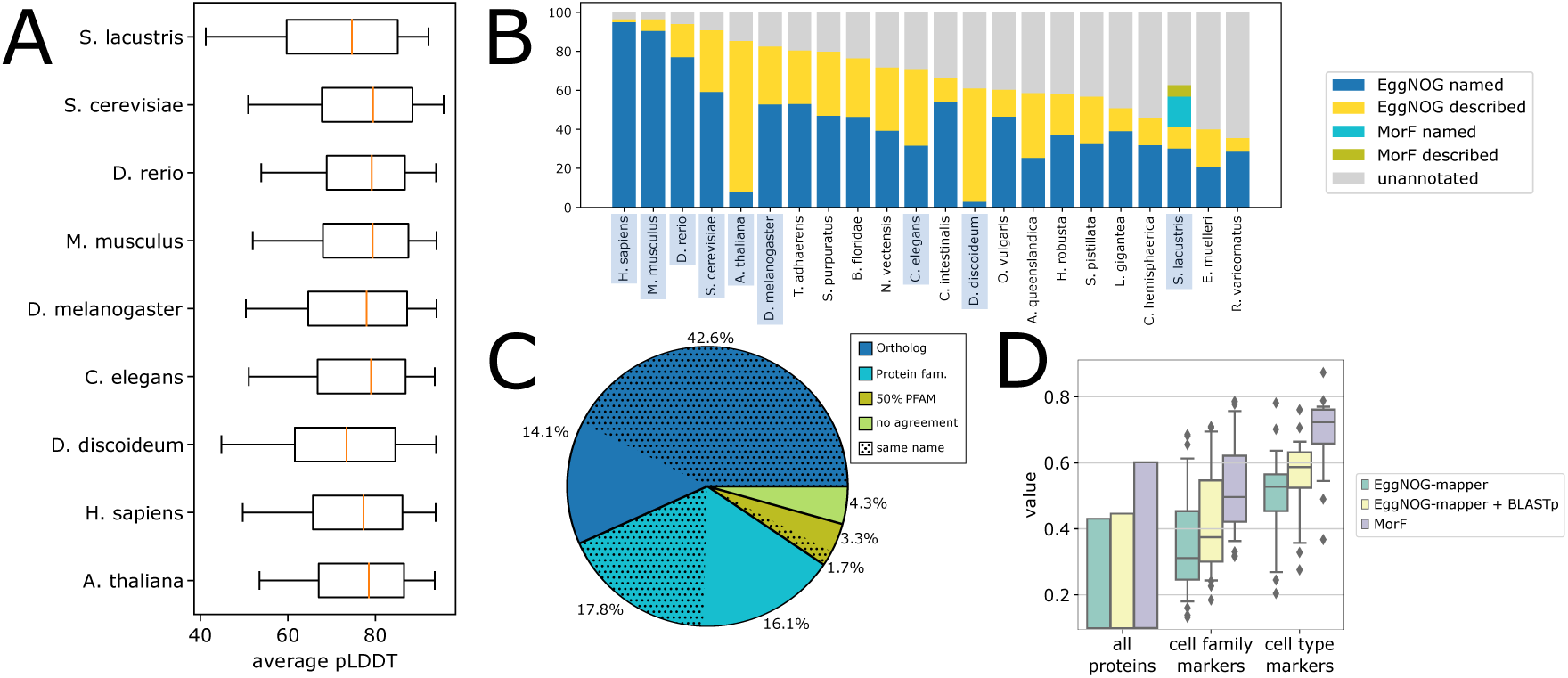
Structural prediction and alignment of the *Spongilla* proteome. A) Distribution of average pLDDT for predicted proteomes from common model species in comparison to *Spongilla lacustris*. B) Proportion of EggNOG or MorF protein annotations in *S. lacustris* and other eukaryotes. Highlighted organisms appear in A. C) Overlap between EggNOG and MorF annotations. *Ortholog* : proteins identified as belonging to the same orthology group in the most recent common ancestor in the EggNOG database. *Protein fam.*: proteins identified as belonging to the same eukaryote orthology group in the EggNOG database, indicating annotations represent homologs in the same gene family. 50% *PFAM* : half of the sequence-based PFAM domains are shared. *No agreement* : MorF and EggNOG annotations identify non-homologous proteins. Subcategories with “same name” denote the fractions where EggNOG and MorF returned the same preferred name for a protein. D) Annotated proportions of different categories of *S. lacustris* proteins.

Next, we used the predicted protein structures as queries to search with Foldseek [8] against AlphaFoldDB [13], SwissProt [14], and PDB [15]. In general, bit scores for the best Foldseek hits were positively correlated with mean pLDDT. Neither parameter correlates with predicted physico-chemical properties of the proteins such as hydrophobicity, isoelectric points or their instability index (Suppl. Note B, Suppl. Figs. S2-S4) [16]. After removing lower-quality matches, we retrieved functional annotations for the best morphologs using EggNOG-mapper (emapper) [17], a state-of-the-art orthology database [18], and then assigned these annotations to the protein in *Spongilla*. This produced annotations for slightly more than 60% of the proteome (25,232 proteins), representing an increase of approximately 50% compared to when *Spongilla* protein sequences were directly searched with emapper. Whereas the usage of emapper is not compulsory for MorF, it provided functional descriptors like EC numbers or GO terms for orthologous groups, facilitating later programmatic comparisons to sequence based methods. However, for downstream biological analysis, gene names and descriptions remain the most succinct, human-readable proxies for protein function. We therefore decided to use the preferred name and description of the best morpholog for each *Spongilla* protein assigned by emapper.

We also compared our results to “legacy” annotations from the recently published *Spongilla* cell type atlas, which used BLASTp to supplement emapper annotations [9]. Compared to this combined sequence-based approach, MorF annotates more proteins proteome-wide (*∼*60% to *∼*40%). More importantly, MorF markedly improved the proportion of annotated cell type and cell family-specific markers (*∼*70%) compared with sequence-based approaches (*∼*56%; Fig. 1D, Suppl. Fig. S7), even considering sequence profiles (Suppl. Fig. S11).

#### MorF annotations agree with sequence-based annotation transfer

Traditional approaches for functional annotation use protein sequences to identify orthologous or homologous proteins. These methods exploit the fact that homology is often the best predictor of shared function, and effectively treat functional annotation synonymously with assessment of homology, although it is known that divergence of function within orthologs is possible [19]. As a next step, we evaluated whether MorF annotations are congruent with those produced via traditional sequence-based homology approaches.

We first examined the agreement between morphologs and homologs on available predicted structures of model organisms. We aligned AlphaFoldDB against itself and kept for each query the top morpholog outside the species taxonomic unit. This ensured that MorF would not be identifying quasi identical proteins from closely related species (e.g. *M. musculus* and *R. norvegicus*), simulating a realistic use case where MorF would be used to annotate a non-model organism without well-studied close relatives (Suppl. Table 3). We assessed performance by comparing the eukaryotic orthologous group of the top morpholog to that of the query protein, as defined by the EggNOG v5.0 database. MorF routinely identifies 75-90% of all available homologs, indicating that it can successfully reproduce sequence-based homology inference across large evolutionary distances.

We proceeded to repeat this analysis with the predicted *Spongilla* structures. A total of 16,589 proteins were annotated by both MorF and emapper. For 90.6% of these proteins the MorF annotation was homologous to the EggNOG assignment (Fig. 1C), being either orthologs (56.7% in the same metazoan orthologous group) or in the same gene family (33.9% in the same orthologous group at the root level). Proteins that share the same gene family but are not annotated as orthologs either represent paralogs or have been misclassified, a problem for orthology inference that is prone to occur with large evolutionary distances [20]. In the remaining 9.4% of cases approximately half shared a majority of their PFAM domains [21]. We explore the overlap between MorF and EggNOG in more detail in Suppl. Note C. Repeating this analysis with sequence profiles produced very similar results (Suppl. Note I).

#### Morphologs share function over long evolutionary distances

As a next step, we sought to explore whether MorF can be used for functional annotation in cases where the evolutionary distance is too large for sequence-based approaches. To test this, we performed Foldseek searches for the predicted *S. cerevisiae* and *A. thaliana* proteomes against *H. sapiens*, and identified morphologs that lacked evidence of homology based on sequence. It is important to note that it is impossible to know for sure whether or not these represent homologs or protein structures that evolved via convergent similarity.

Nevertheless, for the remaining candidates, we tested their functional similarity by examining the overlap of their Enzyme Commission (EC) number where available. The EC number is a four-digit numerical description of enzyme function, with each number representing a progressively finer classification of the enzyme. Agreement on the first digit indicates two proteins are in the same broad enzyme class (oxireductases, hydrolases, ligases, etc.), while complete agreement means that they catalyze the same reaction.

For yeast, 109/145 (75%) enzymes agreed with their human morphologs on three of four EC positions and 53/145 (36.5%) agreed on all four. Similarly, for *Arabidopsis*, 357/532 eligible enzymes agreed to the third EC position (67%) and 176/532 (33%) had identical EC numbers. These results indicate MorF can accurately predict function even in cases where protein homology is unclear due to large evolutionary distances. Furthermore, this is consistent with other work in the field that has demonstrated that structure similarity uncovers homologs between *Homo sapiens* and different *Saccharomyces* species [22] using a similar methodology. We eagerly expect more insights on this topic in the coming months and years.

We next assessed the consistency of the functional annotation for top morphologs in *Spongilla*. For each protein we queried the EC number of all morphologs in the 90th percentile of the Foldseek score range. This serves as an indirect way of validating that significant structural similarity correlates with functional conservation. In the 7072 cases that we could evaluate, the top morphologs were close to identical to the EC number of the best morpholog (average agreement 3.7 positions; see Suppl. Note D).

We also examined consistency between MorF and sequence-based annotations for *Spongilla* by comparing GO term overlap, semantic similarity and depth [23], drawing on the annotation comparisons of the CAFA challenge [24]. Almost all (99.9%) proteins annotated by both strategies have at least some overlap in GO terms with 60.6% being identical and 39.3% partially overlapping to various degrees (Suppl. Fig. S6A). GO term semantic similarities between proteins with overlapping (but non-identical) GO annotations additionally reach an average score of 80 *−* 90% in the molecular function ontology (Suppl. Fig. S6B).

#### MorF expands annotation of signalling pathways in Sponge pinacocytes

A principal goal of functional annotations is to help identify cellular and molecular processes in large-scale genomic datasets. As a next step we used MorF to revisit a recent single-cell RNA-sequencing dataset [9] which allowed us to confirm as well as expand the understanding of cell type functions in sponges. Musser *et al.* showed that sponge pinacocyte express members of the FGF, TGF/BMP, and Ephrin developmental signaling pathways [25, 26]. Pinacocytes are contractile epithelial cells that line the sponge canal system, playing important roles in morphogenesis, barrier formation, and sponge whole-body contractions [27, 28]. Using MorF, we identify morphologs of additional members of these pathways expressed in pinacocytes, extending our understanding of their function in sponges.

In the FGF pathway, previous sequence-based annotations identified Fgf receptors and the FGF regulators Frs and Grb2, which are expressed in different pinacocyte cell types (Figure 2A). Extending this, MorF identified morphologs for GAB1 and GAB2 (GBR2-associated-binding protein 1/2) as well as PTPN11/SHP2, which are necessary for signal transmission into the cell [29, 30]. Notably, MorF also detected a morpholog of the FGF ligand in *Spongilla*, which was not found using sequence-based approaches. Structural superposition of the protein with its best Foldseek hit (UniProtID: P48804, *Gallus gallus* FGF4) revealed an extensive alignment of large parts of the protein, with a RMSD of 0.89 over 543 atoms despite a sequence identity of only 11.8% (Fig. 2B). Newly annotated players in the FGF pathway also exhibited enrichment in pinacocytes, consistent with its previously known cellular role.

**Fig 2.**
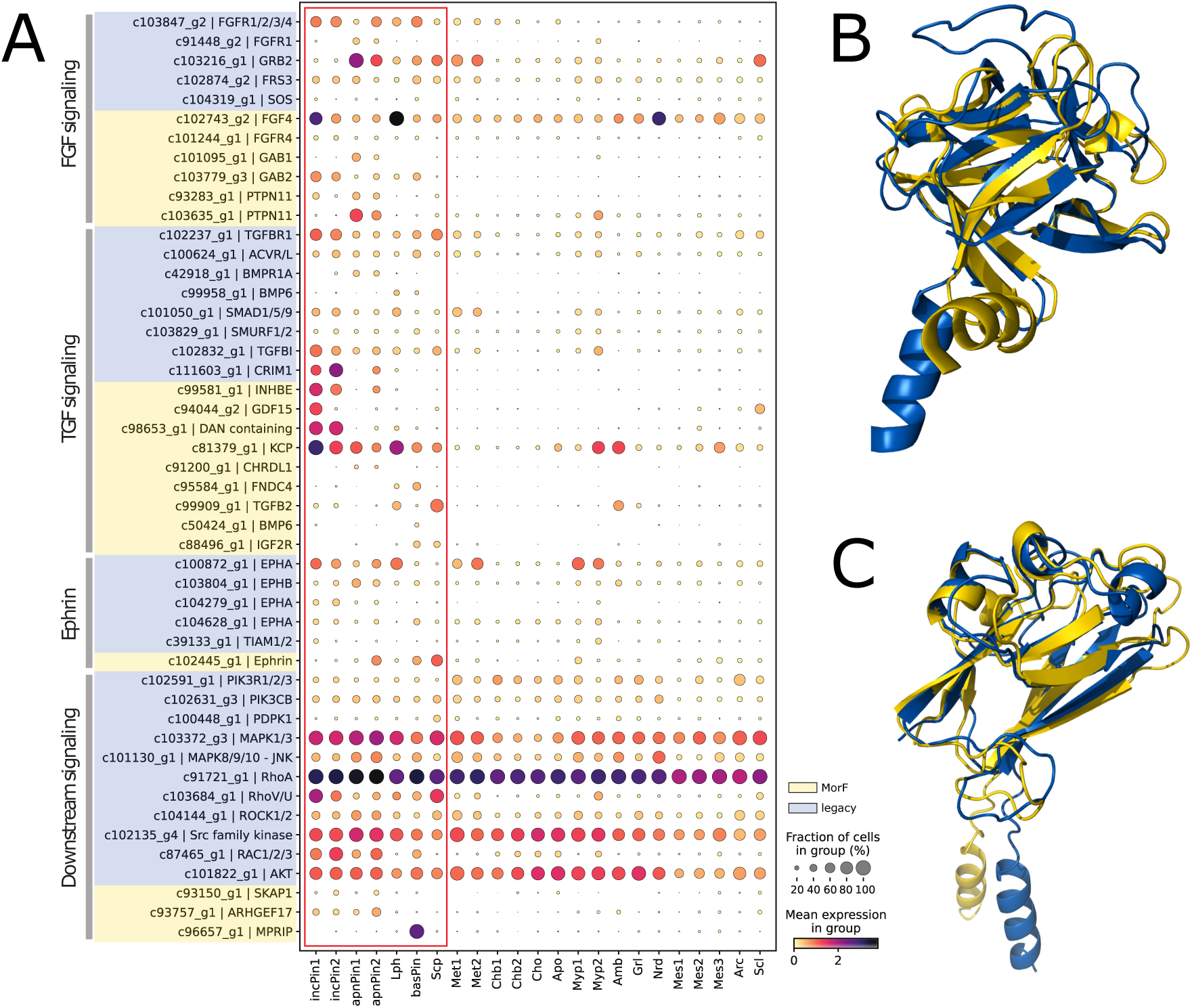
Signalling pathways in *Spongilla* pinacocytes. A) Dotplot of pinacocyte signalling and effector genes. Cell types of the pinacocyte family are encased by a red square. Genes on blue background are annotated by sequence-based methods (“legacy”). Genes on yellow background are annotated by MorF. B) Superposition of *Spongilla* FGF (blue, 61 - 230 aa) and *Gallus gallus* FGF4 (UniProtID: P48804) (yellow, 54 - 194 aa) (RMSD = 0.89 over 543 atoms). C) Superposition of *Spongilla* ephrin (blue, 1 - 153 aa) and *C. elegans* efn-3 (UniProtID: Q19475) (yellow, 29 - 179 aa) (RMSD = 1.59 over 580 atoms). In both cases the structural similarity is apparent despite their low sequence identity of 11.8% and 22% respectively. Superpositions were created using the super command in PyMOL (v2.3.5). Cell type abbreviations: incPin - incurrent pinacocytes; apnPn - apendopinacocytes; lph - lophocytes; basPin - basopinacocytes; scp - sclerophorocytes; met - metabolocytes; chb - choanoblasts; cho - choanocytes; apo - apopylar cells; myp - myopeptidocytes; amb - amoebocytes; grl - granulocytes; nrd - neuroid cells; mes - mesocytes; arc - archaeocytes; scl - sclerocytes.

Musser *et al.* also described multiple genes involved in TGF-*β* signalling in the pinacocyte family, including Tgfbr1, Acvr, Smad and Smurf [31, 32]. MorF extends the list of known actors in pinacocyte TGF signalling, adding morphologs of important ligands such as INHBE, CHRDL1 or KCP.

Lastly, Ephrin and the ephrin receptor (Eph) are membrane-anchored signalling molecules that mediate communication between adjacent cells [33, 34]. Whereas multiple ephrin receptors were detected via sequence similarity, ephrin itself was only found in the *Spongilla* proteome using highly sensitive HMM profile searches [35, 36]. MorF annotates a *Spongilla* gene with differential expression in various pinacocytes as a morpholog of *Caenorhabditis elegans* Efn-3 (UniProtID: Q19475). Although these proteins share only 22% sequence identity, the superimposed structures achieve an RMSD of 1.59 over 580 atoms (Fig. 2C). A separate Hmmer search [37] using the ephrin Pfam profile (PF00812) picked up the same gene, supporting MorF to be at least as sensitive as curated HMM profile searches (also see Suppl. Note I).

FGF, TGF-beta as well as Ephrin interestingly exhibit converging downstream signalling pathways, including PI3K/Akt, ERK/MAPK1, JNK or RhoA/ROCK, responsible for cell growth, differentiation, migration and cytoskeletal organisation [33, 38, 39]. Together, MorF and sequence-based methods identified morphologs of principal proteins involved in the downstream pathways. The genes encoding these proteins are broadly expressed across most *Spongilla* cell types, consistent with their diverse functional roles. This example highlights the power of MorF to vastly extend annotations and further elaborate cell type specific functions.

#### Redox metabolism and control in myopeptidocytes

The mesohyl of sponges is a collagenous, dynamic tissue forming large parts of the body between pinacocytes and the feeding choanocyte chambers [40]. Musser *et al.* identified five novel mesenchymal cell types [9] in *Spongilla*. Among them, myopeptidocytes are an abundant uncharacterised cell type, forming long projections that contact other cells. Sequence-based annotations suggested myopeptidocytes function to generate and degrade hydrogen peroxide by expressing dual oxidase (Duox1), its maturation factor (DuoxA), and Catalase (Cat) [41].

Myopeptidocytes also express transporters of copper ions as well as Ferric-chelate reductase (FRRS1 and FRRS1L), which recycle Fe^3+^ to its reduced state, and suggests iron-based generation of H_2_O_2_. In their reduced state metal ions react with H_2_O_2_ (Fenton reaction) [42] leading to the generation of hydroxyl radicals in cells. The existence of these prominent reactive oxygen species (ROS) (Fig. 3) is further supported by expression of Cyba [43]. However, further roles of ROS metabolism and function are unclear.

**Fig 3.**
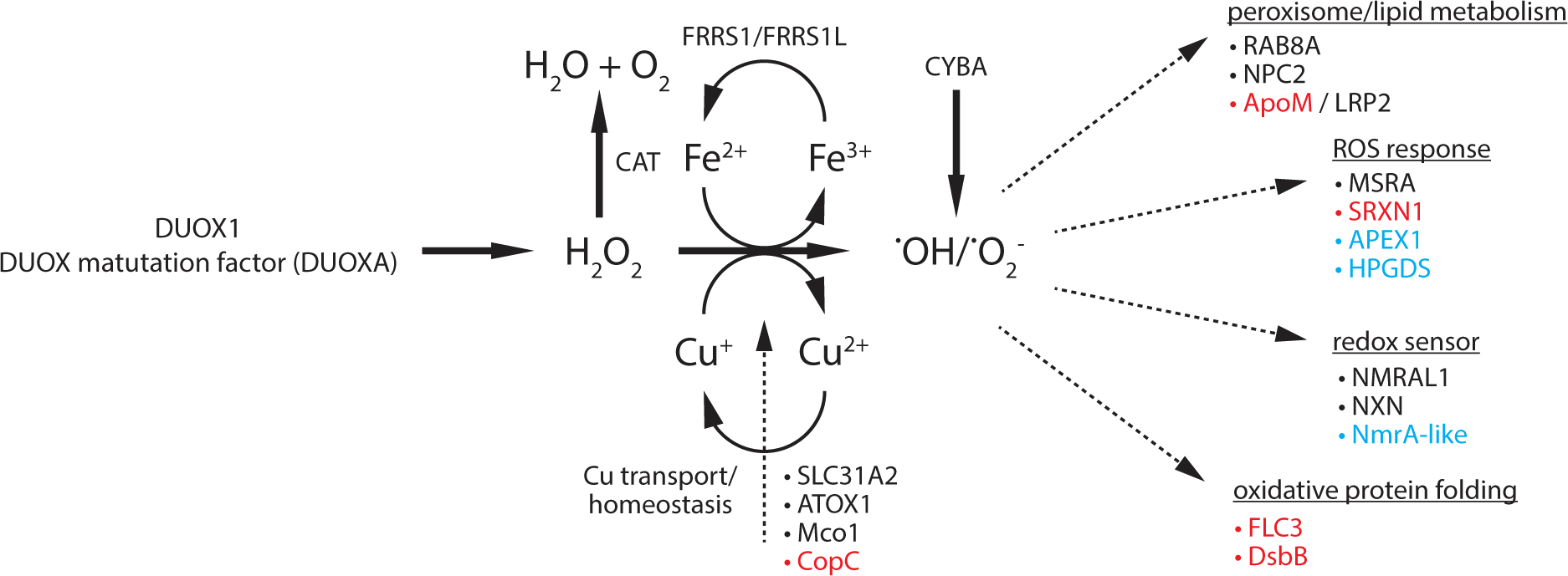
ROS metabolism and redox-control in Myopeptidocytes. Myopeptidocytes differentially express multiple genes involved in redox control and ROS defence. Genes in black have been annotated using sequence based methods. Blue proteins have protein family level sequence based annotation with updated functions inferred by MorF. Genes in red have been functionally annotated using MorF.

MorF predicted the functions of key additional members of ROS generation, metabolism and response that are expressed in myopeptidocytes (Fig. 3, Suppl. Table 5). Morphologs of disulfide oxidoreductase (DsbB) as well as Flavin carrier protein (FLC), both playing a role in oxidative protein folding, have been detected by MorF. [44, 45]. NmrA-like proteins act as redox sensors in the cell [46]. Consistent with a possible redox regulation role, myopeptidocytes express morphologs of a range of additional ROS responsive proteins: Sulfiredoxin 1 (SRXN1) promotes resistance against oxidative stress damage [47], whereas AP endonuclease 1 (APEX1) protects against ROS induced DNA damage [48, 49]. We also identified a glutathione S-transferase member (HPGDS) which together with previously annotated methionine sulfoxide reductase (msrA) is an important enzyme involved in the repair of proteins damaged by oxidative stress [50, 51]. Finally, myopeptidocytes express morphologs of the LDL-receptor LRP2 and its binding partner apolipoprotein M (ApoM) [52], suggesting a role in lipid metabolism and consistent with the observation myopeptidocytes exhibit round inclusions that may represent lipid droplets. Interestingly, lipid metabolism and redox control is tightly coupled in peroxisomes which are responsible for beta-oxidation of long-chain fatty acid [53]. Structure based annotation of myopeptidocyte marker genes enabled us to substantially hypothesise about the role of this unexplored cell type in sponges.

#### Polysaccharide hydrolysis in enigmatic mesocytes

Mesocytes are newly discovered medium-sized sponge cells whose name refers to their location in the mesenchymal mesohyl [9]. The single-cell RNA-seq data produced a series of marker genes specifically expressed in the mesocyte cell clusters; however the lack of annotation for many of these genes made it difficult to hypothesise functions for these cell types.

New annotations provided by MorF include proteins such as expansin (yoaJ), glucan endo-1,3-beta-glucosidase (BG3), and spore cortex-lytic enzyme (sleB), all hydrolases that specifically degrade cell walls, cellulase, chitin, other polysaccharides [54–57] and proteins (Table 1, Suppl. Table 6, Suppl. Fig. S8). It is thus tempting to speculate that mesocytes represent cells specialised to digest polysaccharides that are otherwise difficult to hydrolyse. For instance, the mesohyl has been shown to contain chitin which likely helps provide structural support to the sponge body [58]. The presence of chitinase in mesocytes suggests a possible role as structural remodelers of the sponge endoskeleton.

**Table 1.**
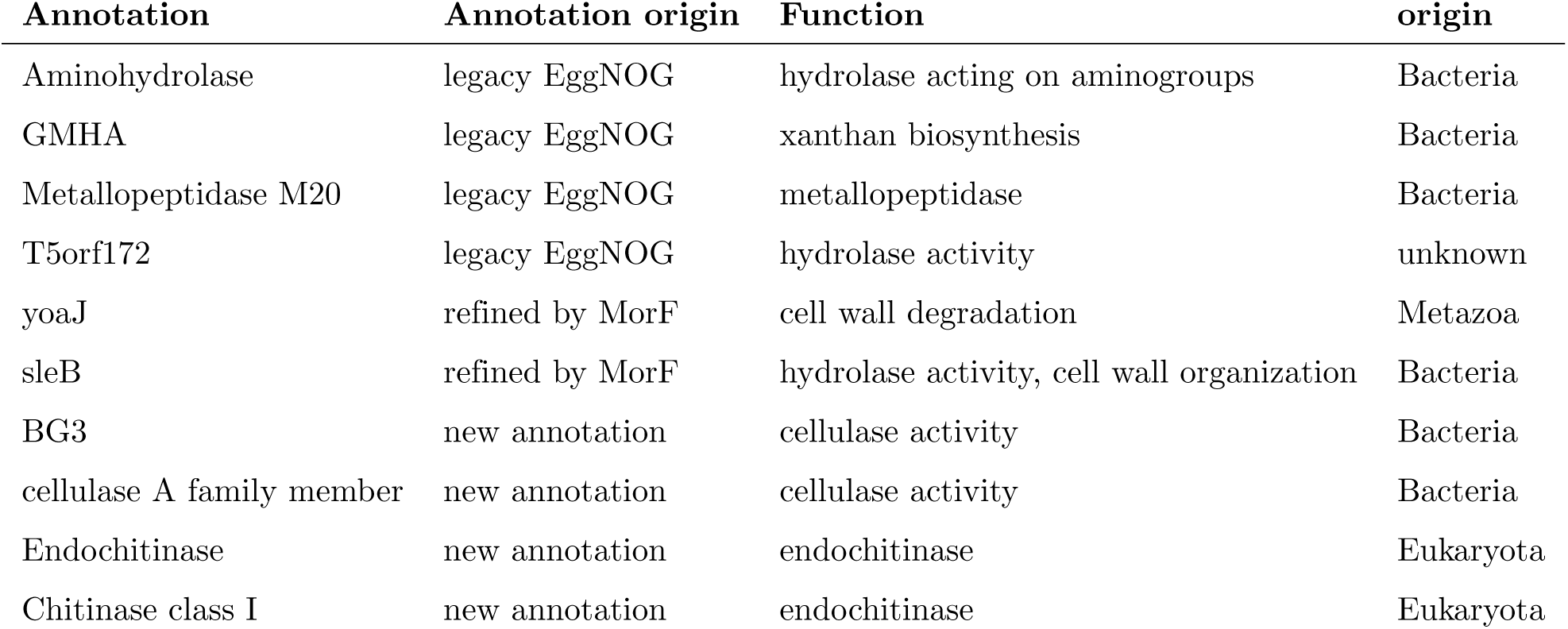
Hydrolytic enzymes of Spongilla mesocyte marker genes.

The sponge mesohyl contains digestive and phagocytic cells [59] that process food particles captured by pinacocytes and choanocytes. These food particles often include bacteria and algae, which are protected by polysaccharide and glycoprotein cell walls [60] that require specialised enzymes to break down. While those enzymes are notably absent from the metazoan digestive repertoire [54], sponges are at least demonstrably able to digest algae [59].

The absence of these mesocyte-specific hydrolytic enzymes from the digestive toolkit of animals suggests four possibilities for their appearance in the *Spongilla* single-cell data: that they are an artefact (contamination), that they are an evolutionary novelty within *Porifera*, that they were lost in all other animal lineages, or that they were acquired via horizontal gene transfer (HGT).

To explore these different possibilities, we used the marker gene sequences to find putative homologs in the RefSeq non-redundant (*nr*) [61] and metagenomic databases. Strikingly, the best hits were mostly of bacterial origin, exhibiting 40 *−* 70% shared sequence identity with sponge proteins, however mostly lacked annotation (Suppl. Table 1). Notably, we identified putative homologs for each gene in other sponges, suggesting the presence of these sequences in the *Spongilla* protome is unlikely to have occurred due to contamination (Fig. 4B). Consistent with this, we found codon usage and GC content for these genes did not deviate from the *Spongilla* background (Fig. 4A). Lastly, we located all candidate genes on different long contigs (avg. length *∼*420kb) of an in-house draft assembly of the *Spongilla* genome. The specific co-expression of functionally similar proteins in mesocytes is in contrast to a random contamination.

**Fig 4.**
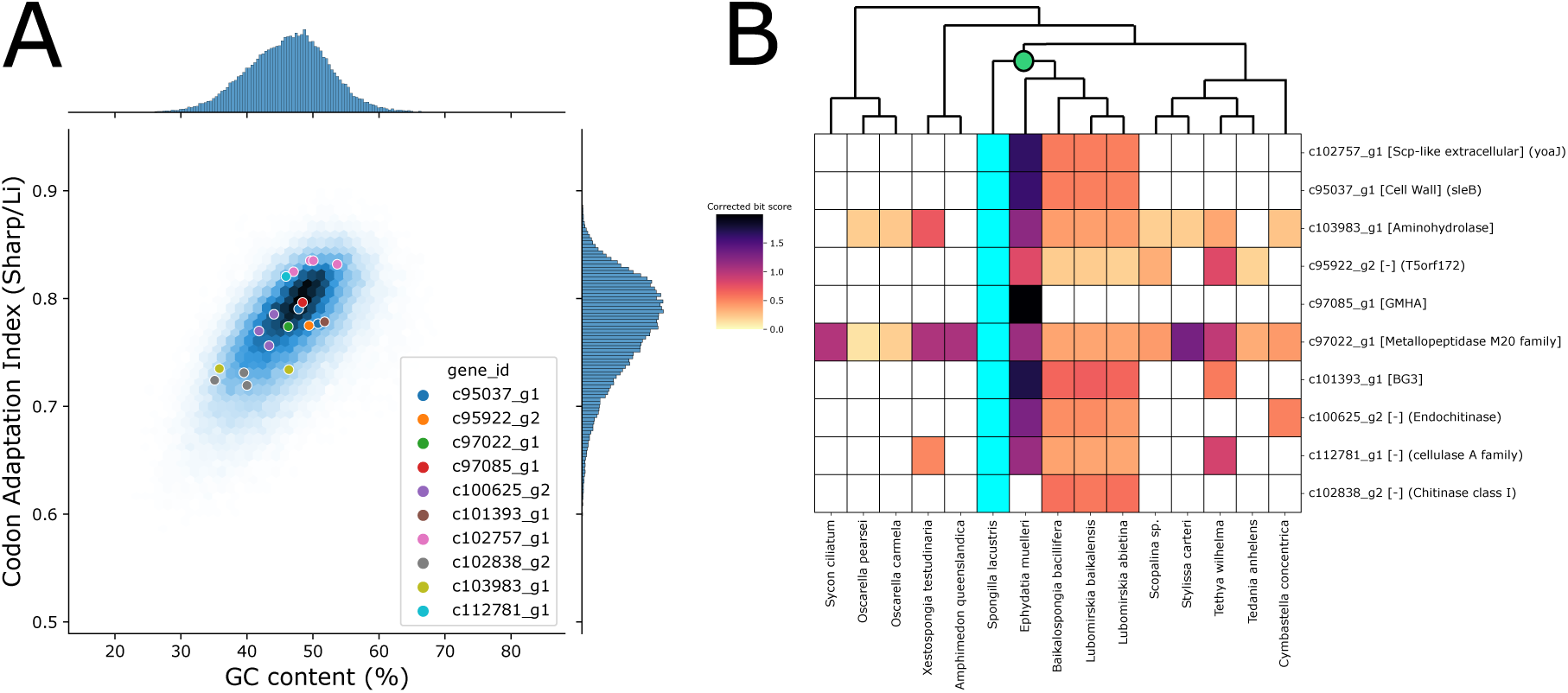
Horizontal gene transfer of mesocyte marker genes. A) Distribution of GC-content and codon usage of mesocyte marker genes of non-metazoan origin (colored dots) compared to the entire *Spongilla* transcriptome (blue background). B) Heatmap showing scores of best search hit of *Spongilla* mesocyte marker genes in various sponge species. The green dot denotes the last common freshwater sponge ancestor.

The prospect of HGT is tantalising. Proteins with enzymatic functions like the ones in the *Spongilla* candidates (polysaccharide hydrolases and metallopeptidases) have been proposed to be horizontally transferred in *A. queenslandica*, a marine demosponge, *S. rosetta*, a choanoflagellate, and *M. leidyi*, a ctenophore [62–64]. Additionally, *Spongilla* genes c97022 g1 and c103983 g1, a putative aminohydrolase and metallopeptidase respectively, are not only broadly distributed within sponges, but can also be found in the proteomes of choanoflagellates *S. rosetta* and *M. brevicollis* (Suppl. Note L). Furthermore, the *S. rosetta* targets with highest similarity to the *Spongilla* sequences had already been identified as horizontally transferred genes [63]. This would tentatively place this HGT event at least before the split of choanoflagellates and animals (more than *∼*500mya). Similarly, the phylogenetic distribution of c102757 g1, c95037 g1, and c102838 g2 (yoaJ-sleB-Chitinase class I) would indicate that this group of genes was acquired with the colonisation of freshwater environments (*∼*15 *−* 300mya).

Although our analyses suggest these genes may have originated via HGT [65], it is important to consider alternative explanations. One possibility is that these genes are the results of widespread bacterial contamination in sponges genomes and proteomes. Other possibilities include the repeated loss of these genes in other animal lineages, or the convergent evolution of similarity with bacterial proteins. Additional confirmation of their presence in sponge genomes, or evidence of RNA transcripts in sponge cell nuclei, would help validate the hypothesis they arose via HGT. Regardless of the source of the genes, MorF annotations provided a novel hypothesis for the elusive function of sponge mesocytes, helping uncover new aspects of sponge biology.

#### Identifying novel well-folded proteins in *Spongilla lacustris*

Using the MorF pipeline with a stringent bit score cut-off and augmenting it with sequence-based annotations we annotated a total of 26,633 out of 41,943 predicted proteins. The remaining 15,312 proteins may represent incomplete fragments, untranslated sequences, or sponge lineage-specific genes. Notably, we found 3,875 unannotated proteins with a pLDDT score greater than 70, indicating well-folded structures. Although many of these had poor Foldseek alignments falling below our accepted bit score threshold, 316 had no Foldseek hit whatsoever. Manual inspection revealed that the overwhelming majority of these are predicted to be long helices, except for 35 non-helical structures. To ensure these hadn’t somehow eluded sequence similarity searches, we used NCBI BLASTp to identify potential homologs in the NR database, even very remote ones (Suppl. Table 2). Seven of the sequences find no matches at any theshold nor get significant PFAM domain hits. Notably, several of them are broadly expressed in *Spongilla* cell types, presenting prime candidates for truly novel structures.

Recent changes in the amount of available structures are almost certainly going to affect these numbers. The latest AlphaFoldDB version contains predicted structures for more than 214 million proteins [66], and with the continuous deposition of more sequences and their predicted structures our view of protein structure space will be ever closer to complete.

### Discussion

We predicted the protein structures of the entire proteome of the freshwater sponge *Spongilla lacustris* and aligned them against model organism proteomes, PDB, and SwissProt. This approach increased the annotation of the proteome from 40% to 60%, an approximately 50% increase. We found that in more than 90% of cases, sequence-based and structure-based annotations identified homologous proteins, a finding supported by recent work with well-annotated model species [22]. Additionally, these proteins overlap largely in their corresponding GO terms and their semantic similarity.

In *∼*5% of cases, MorF and sequence-based approaches identify unrelated proteins. These may be the result of technical artifacts (Suppl. Note C), but may also indicate example of protein structures evolved similarity to unrelated proteins via evolutionary convergence. In the future, it will be interesting to explore these cases experimentally to test whether structural similarity or homology is a better predictor of protein function [67].

To demonstrate the usefulness of MorF for functional annotation, we revisited the cell type marker genes previously identified from *Spongilla* scRNA-seq data [9]. In the epithelial pinacocyte family we significantly improved annotation by detecting morphologs of key players in FGF, TGF-*β*, as well as ephrin signalling. We were able to infer complex modes of redox regulation as a possible function of the myopeptidocytes. Finally, we detected polysaccharide and protein hydrolyzing enzymes potentially used for digesting cell walls in the so far enigmatic mesocytes, which may have originated from bacteria via HGT. If proven to be true, this example reveals cell types whose functional role is largely defined by genes acquired via HGT. A comparable example has been found in nematodes, which expanded their diet after the HGT of cellulase from a non-animal eukaryote [68]. These events are remarkable because they suggest HGT may help give rise to new cell type-specific functions [69]. Moreover, the large evolutionary distance between animals and bacteria has very likely hindered the identification of these events, and we anticipate MorF will uncover additional examples in other species. Recent evidence from nematodes showed an expansion of dietary possibilities after HGT of a eukaryotic cellulase gene. By re-evaluating cell type marker genes from the *Spongilla* cell type atlas it became obvious that structure-based annotations not only expand the molecular context of known cell type functions but also allow substantiated hypotheses about previously unexplored cell types. These hypotheses can serve as a next step for further investigation and discoveries.

Well-folded *Spongilla* proteins without any annotation constitute intriguing candidates for novel protein folds and functions specific to sponges. Many of these proteins are predicted to be long alpha helices. This could either be an artifact of spuriously translated proteins (as described in [70]) or present sponge specific proteins for structural integrity of the sponge body, similar to the “constant force spring” character of naturally occurring single *α*-helices [71].

However, unannotated proteins with globular structures, expressed in the scRNA-seq data, are most probably functional and offer a great resource to study the evolutionary emergence of protein folds.

## Conclusions

The lack of reliable functional annotation has so far been a major bottleneck in the analysis of -omics datasets from non-model species, in particular those separated from traditional models by large phylogenetic distances. Here, we demonstrate that by exploiting the evolutionary conservation of protein structure it is possible to dramatically improve protein functional annotations in non-model species. We show that proteins with significantly similar structures (morphologs) are often homologs. Using GO-terms as well as EC numbers as measures for functional similarity, we illustrate that in many cases morphologs are functionally similar across large evolutionary distances and can therefore be used for functional transfer. Although protein structural predictions for an entire proteome might be outside the technical capabilities of many labs, the pipeline described here can be used to query individual highly informative candidate genes from proteomics or single-cell -omics experiments. During the peer review process of this manuscript, protein structures for the entire UniProt database were predicted and updated version of Foldseek (v4) as well as the EggNOG database (v6.0) have been released. It is reasonable to expect that in the future protein sequences deposited in public databases will automatically receive predicted structures, paving the way for unique insights into biological functions across the tree of life.

## Materials and methods

In the manuscript, “annotation” of a protein refers to the existence of an emapper based preferred gene name or description. In all boxplots, the box extends from the lower to upper quartile values of the data, with a line at the median. The whiskers represent the 5 *−* 95% percentiles.

### Sequence-based annotation of the *Spongilla lacustris* proteome

Juvenile freshwater sponges (*Spongilla lacustris*), grown from gemmules, were used for bulk RNA isolation and sequencing. *De novo* transcriptome assembly with Trinity, returned 62,180 putative isoforms, covering 95.2% of Metazoan BUSCOs [72]. To identify putative proteins, Transdecoder [73] (version 3.0.1) was used with a minimum open reading frame length of 70 amino acids, resulting in 41,945 putative proteins. The longest putative protein per gene ID was kept. The resulting predicted proteome was annotated by EggNOG mapper [17, 18] (v2.1.7, default settings) via the website.

### Legacy annotation

Musser *et al.* [9] used the putative proteins to create a phylome by constructing gene/protein trees for each protein [74]. The phylome information was used to refine the assignment of transcripts to genes. In some cases, 3’ and 5’ fragments of a gene were assigned to two different transcripts. These fragments were merged into the same merged gene name using the gene tree information. Functional annotations were supplemented by EggNOG mapper (v1) and blastp searches against human RefSeq (default parameters). This annotation was used in the original *Spongilla* scRNA sequencing publication and is present in the single-cell data. The legacy annotation was used for the single-cell data analysis, but not for the comparison between the MorF pipeline and the sequence-based annotation transfer.

### Structure-based annotation of the *Spongilla lacustris* proteome

#### MorF and constituent tools

We designed a simple pipeline that goes from sequence to predicted structure to functional annotation (symbolically, **Mor**pholog**F**inder, or MorF), and used it to annotate the *Spongilla lacustris* predicted proteome. We used ColabFold to predict structures and subsequently aligned the predicted structures against all currently available (solved and predicted) protein structures using Foldseek. Finally, we used structural similarity to transfer annotations from the morphologs to their corresponding *Spongilla* protein queries. In the following we show an overview of the tools in use and a more detailed description of the MorF pipeline.

**MMseqs2** (version 92deb92fb46583b4c68932111303d12dfa121364) [75] is a software suite for sequence-sequence and sequence-profile search and clustering. It is orders of magnitude faster than BLAST at the same sensitivity and is widely adopted [17].

**AlphaFold2** [7] is a neural network-based model that predicts protein three-dimensional structures from sequence, regularly achieving atomic accuracy even in cases where no similar structure is known. AlphaFold is widely considered to have revolutionised the field of structural bioinformatics, greatly outperforming the state of the art in the most recent iteration of the CASP challenge [76].

AlphaFold quantifies prediction confidence by pLDDT, the predicted local distance difference test on the *Cα* atoms. Regions with pLDDT *>* 90 are modelled to high accuracy; regions with 70 *<* pLDDT *<* 90 should have a generally good backbone prediction; regions with 50 *<* pLDDT *<* 70 are low confidence and should be treated with caution; regions with pLDDT *<* 50 should not be interpreted and probably are disordered.

**ColabFold** [10] is a pipeline that combines fast homology searches via MMseqs2 [75] with AlphaFold2 [7] to predict protein structures 40 to 60 times faster than the original AlphaFold2. We installed colabfold locally from the localcolabfold repository [77], version 1.4.0 **Foldseek** [8] enables fast and sensitive comparison of large structure databases. Foldseek’s key innovation lies in the appropriate translation of structure states to a small alphabet, thus gaining access to all the heuristics of sequence search algorithms. We used version 3-915ef7d.

#### Structural databases

We used the **AlphaFold database** v1 [13], containing over 360,000 predicted structures from 21 model-organism proteomes, as provided by Foldseek v3-915ef7d.

From Foldseek v3-915ef7d we also used the **Swiss-Prot** [6] and **PDB** [15] databases.

#### The MorF pipeline

A visual representation of the MorF pipeline can be found in supplement figure S1. To predict structures, we adapted the ColabFold pipeline as outlined in [78].

##### Multiple sequence alignment generation

We downloaded reference sequence databases (UniRef30, ColabFold DB) and calculated indices locally ([79, 80], adapted from ColabFold setup databases.sh). We were interested in homology detection at the limit of the twilight zone, so UniRef30, a 30% sequence identity clustered database based on UniRef100 [81], was the adequate choice. We calculated MSAs for each *Spongilla* predicted protein ([82], adapted from [83]) using MMseqs2 [75].

##### Structure prediction

We predicted structures for all *Spongilla* predicted proteins using ColabFold [10] as a wrapper around AlphaFold2.

We split the MSAs in 32 batches and submitted each one to the EMBL cluster system (managed by slurm [84]); we used default arguments but added --stop-at-score 85 [85]. The calculations were done on NVIDIA A100 GPUs, on computers running CentOS Linux 7. We used GCC [86] version 10.2.0 and CUDA version 11.1.1-GCC-10.2.0 [87]. We processed the resulting PDB-formatted model files with Biopython’s PDB module [88].

##### Structure search and annotation transfer

Structural search was conducted using Foldseek which allows fast comparison of large structural databases. We downloaded PDB, SwissProt, and AlphaFold DB. For each *Spongilla* protein we kept the best-scoring AlphaFold2 model, and used them to construct a Foldseek database. These models were then used to search against the three structural databases (see [89]). For each search we kept the Foldseek hit with the highest corrected bit score in each database and aggregated the three result tables (AlphaFoldDB, PDB, SwissProt) into one. We imposed a bit score cutoff of *e*^5^ on Foldseek hits based on their bimodal distribution [90] and personal communication with the Foldseek authors. Annotations of the best hits (= morphologs) were gathered from either UniProt via its API [91] or through EggNOG mapper (v2.1.7, default settings) [17] by using the sequences of the morphologs (pulled by UPIMAPI [92]). To facilitate downstream analysis, we extracted summary tables from each resource type. This procedure can be found in the corresponding notebook [93]. A total of 1401 proteins received sequence annotation but their Foldseek best hits were below the bit score cutoff.

#### Instructions for MorF searches of single proteins using openly available web servers

MorF searches for whole proteomes require large computational resources. However, searches can be carried out for a small number of proteins of interest (e.g. top differentially expressed genes in RNAseq or proteomics experiments) using openly available web tools:

- Prediction of protein structure using ColabFold [10]: The structures of proteins of interest can be predicted using the ColabFold Google Colaboratory notebook [94]. Detailed instructions are described in the notebook. For a quick default run, users paste a protein sequence into *“query sequence”* and hit *“Runtime” - “Run all”*. The results can be downloaded as a zip archive which includes the pdb models of different structure model quality ranks.
- Structure similarity search using Foldseek [8]: The best ranking model (*“…rank 1 model X.pdb”*) can be queried using the Foldseek webserver [95]. Users can upload the pdb file of the model and select databases to use for the search. In default mode, all available databases will be searched. The Foldseek output is structured blast-like and sorted according to best scoring morphologs within the selected databases.
- (Optional) Additional annotation using EggNOG [17]: In order to retrieve additional functional as well as phylogenetic information about the best scoring morpholog, the EggNOG database can be searched [96]. Both protein sequence, as well as UniProt ID can be used to retrieve information about orthologous groups as well as GO-terms, EC numbers, etc.

### GO term semantic similarity and depth calculation

For an in depth comparison of sequence and structure based annotations, we calculated the semantic similarity and depth of GO terms between annotation pairs [23]. Calculation of semantic similarities was done with the stand-alone version of GOGO [97] in default mode using the Average-Best-Match (ABM) method for calculating gene functional similarity [98, 99]. GO term semantic similarity was compared between all GO terms from annotation pairs with partially overlapping GO terms. Calculation of GO term depths was done with the GOATOOLS Python library [100]. GO term depths were compared using the overlapping GO term assignments between sequence- and structure based annotations with partially overlapping GO terms [23].

### Differential gene expression in single-cell transcriptomics data

We obtained the processed Seurat file from [9], and downloaded the lists of differentially expressed genes of clusters, cell types, and cell type clades from the supplemental material of the same publication (Suppl. Data S1 to S3; file science.abj2949 data s1.xlsx; tabs “Diff. exp. 42 clusters”, “Diff. Exp. cell types”, “Cell type clade genes (OU tests)”. The single-cell data operates on the level of genes, so we transformed the sequence-derived and MorF annotations by merging isoform entries and keeping the entry with the best bit score.

We used the legacy annotation included in the file (phylome, emapper, and BLASTp-based), the sequence-derived annotation, and the MorF output to propose names for *Spongilla* proteins. We prioritised sequence-derived annotations (legacy annotation, EggNOG preferred name, EggNOG description) and fell back to MorF (MorF preferred name, MorF description) when there were none. We produced dotplots for the top 200 differentially expressed genes in each cell type and manually inspected them. We focused on terminally differentiated (named) cell types; owing to the presence of continuously differentiating stem cells, the single-cell data contains many clusters of maturing or differentiating cells whose expression patterns do not distinguish them from their mature counterparts. Code and detailed explanations are available in the corresponding notebook [101].

### Protein structure visualisation and superposition

In order to visualise predicted structures from the *Spongilla* proteome, we used PyMOL Version 2.3.5 [102]. Superposition with their respective best Foldseek hit was carried out using the super command. super creates sequence- independent superpositions and is more reliable for protein pairs with low sequence similarity.

### Detection of ephrin orthologs in *Spongilla* using HMMER

To validate the ephrin ortholog detected by MorF in *Spongilla*, we recapitulated a previous effort to detect ephrins in different species using extensive HMMER protein profile search [36]. For this, we performed a HMMER search (v3.3.2) using the ephrin Pfam profile (PF00812) against the *Spongilla* proteome using default settings.

### Detecting HGT in Spongilla

To detect HGT we followed the proposal by Degnan [65] and sought to get multiple indications for HGT. In particular: phylogenetic evidence that the candidate gene is more closely related to foreign than to animal genes; genome data showing the candidate gene assembles into a contiguous stretch of DNA with neighboring genes unambiguously of animal origin (this requires, of course, the availability of a sequenced and assembled animal genome; the more complete the assembly, the more confident the HGT identification); and gene sequence revealing metazoan-like compositional traits, including presence of introns, GC content and codon usage. Where possible, gene expression data showing active transcription of candidate genes in animal cell nuclei can enormously strengthen a case, and also addresses the issue of whether or not the HGT-acquired gene is active in its new genomic context.

All putative HGT proteins were used as an input for default blastp searches against non-redundant protein (nr) as well as metagenome databases (env nr). Additionally, default EggNOG 5.0 sequence searches were run. We used SpongeBase [103] to obtain transcriptomes and genomes for 13 sponge species, and obtained a 14th one directly from its repository [104]. We built sequence databases with MMseqs2 version 12-113e3 and searched against them (mmseqs easy-search) with the protein sequences of all isoforms of the HGT candidates. The resulting alignments were filtered to keep the best-scoring alignment per species per gene.

The table of best hits per genome was visualised in Python, and a sponge phylogeny was added manually based on [105] (also A. Riesgo, personal communication).

## Acknowledgements

The authors thank Alexandros Pittis for his technical feedback; the Arendt lab for discussions; the ModelArchive team for their support in publishing the predicted structures; Ana Riesgo for sharing demosponge phylogenies; Juan Daniel Montenegro Cabrera for suggesting the term “morpholog”.

## Availability of data and materials

The dataset supporting the conclusions of this article is available in the Zenodo repository, https://zenodo.org/record/7583930 under DOI 10.5281/zenodo.7583930. The predicted *Spongilla* protein structures are deposited in ModelArchive under https://www.modelarchive.org/doi/10.5452/ma-yoep2.

The code that produced the analysis as well as the figures, including a notebook with instructions on how to obtain the data, is available online. In particular:

- Project name: MorF
- Project home page: https://git.embl.de/grp-arendt/MorF/
- Archived version: v1.1 https://git.embl.de/grp-arendt/MorF/-/tags/v1.1
- Operating system(s): set-up depends on a UNIX environment; analysis is platform-independent
- Programming language: bash, Python, Perl
- Other requirements: for an exactly reproduced environment refer to scripts, conda environment description.
- License: GPL 3.0

## Ethics approval and consent to participate

Not applicable.

## Consent for publication

The publication contains no personal data in any form.

## Authors’ contributions

NP and FR conceived the project. NP, FR, JMM, MM and MS designed the project. NP and FR performed the analysis. MS and MM performed additional sequence-based analysis. NP, FR, JMM, DA wrote the manuscript. All authors read and approved the final manuscript.

## A MorF overview

**Fig S1.**
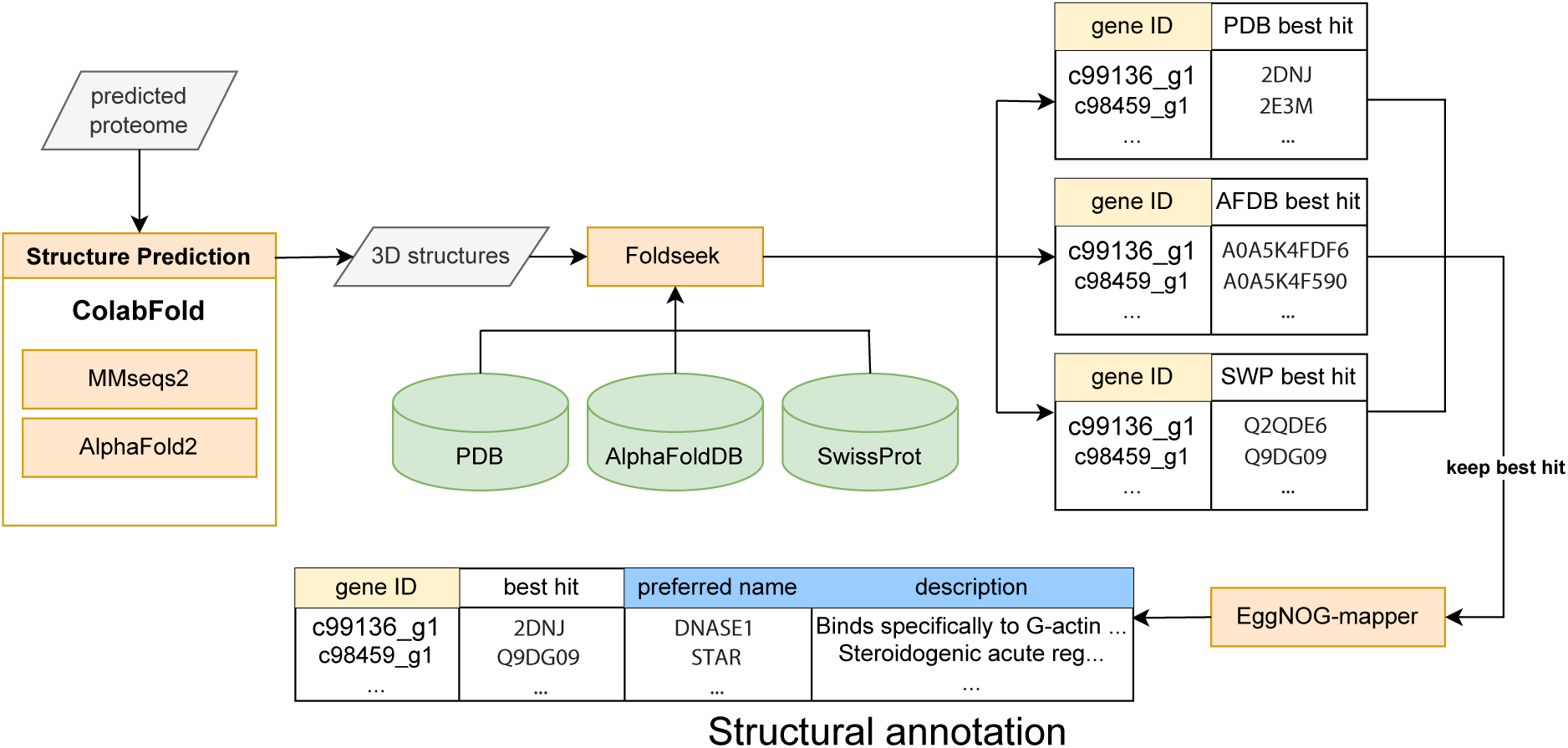
Overview of the MorF pipeline. ColabFold is called with a proteome as input. It produces multiple sequence alignments for each protein using MMseqs2 and predicts structures for each structure using AlphaFold2. The best model per protein is kept and used as query in Foldseek to search against PDB, AlphaFoldDB, and SwissProt. This creates an m8-formatted table for each database, where the significant hits for each query protein are kept. The tables are merged by keeping the target with the highest bit score (= highest scoring morpholog). The sequence of the morpholog is then retrieved from UniProt and used as a query for emapper. This finds the target sequence itself and retrieves the EggNOG entry, including preferred name and description.

## B Extracting quality indicators from the MorF pipeline and comparing against physico-chemical parameters of proteins

The latest Foldseek version changes results are ranked and e-values are calculated [106]. This change came too late in the manuscript revision process for us to repeat all analysis, but we encourage readers to use the latest version. This section refers to the version of Foldseek used in the manuscript (3-915ef7d) and may not be accurate any more.

To get a better understanding of the relationship between the various parameters of the MorF pipeline as well as physico-chemical parameters of the proteins, we assessed the correlation between them (Fig. S2). We hoped to find a combination of measures that together would be a robust indicator of prediction quality for MorF. Additionally we sought to explore the relationship between structural prediction quality and various intrinsic protein parameters.

Most obviously, query length, (uncorrected) structural bit score, sequence bit score (emapper) as well as molecular weight correlate very strongly. Longer proteins (= heavier proteins) will have more or longer aligned regions with their targets, and as the bit score is additive, will accumulate higher bit scores, regardless of whether the alignment happens in structure or sequence space. It seems these measures can be used interchangeably; out of principle, we would prefer to use query length, as it is the least derived.

A way to overcome the dependence of bit scores on the length of the query protein is to normalise by the length of the aligned region (*corrected bit score (FS)*). This measure correlates with the percent structure state identity. This is equivalent to percent sequence identity correlating strongly with the bit score in a sequence similarity search, and is not surprising at all. More interestingly, these measures, together with the MSA size, correlate with the average pLDDT as a measure for protein prediction quality. Nevertheless, this correlation is less pronounced, suggesting to look at pLDDT independently.

Relative alignment length (FS) (percentage of the query (*Spongilla*) structure aligned with the best morpholog) only correlates weakly with other parameters.

Three things become apparent: first, higher pLDDT scores generally correlate with higher Foldseek bit scores. This means that Foldseek does a better job of identifying structural similarity for well-folded proteins. This seems intuitive: the atomic coordinates of well-folded proteins will be more deterministic than disordered proteins, so well-folded orthologs will overlap structurally over their entire length, while heavily disordered orthologs might only overlap structurally over very short patches. Additionally, high pLDDTs are reached when sufficiently large MSAs are available. Large MSAs contribute more information to AlphaFold’s Evoformer module, leading to a higher predictive power.

Second, the relationship between pLDDT and (corrected) Foldseek bit score is not dependent on the length/weight of the protein. Independent of their size, proteins structures are predicted equally good or bad.

Third, pLDDT as well as Foldseek bit score do not correlate with various physico-chemical parameters of the proteins. Protein structure prediction seems to be independent from gravy score [110], aromaticity [109], isoelectric point, instability index [107] as well as the average of flexibility index [108] of proteins. Consequently, Foldseek bit score is not dependent on these parameters.

Next, we were interested in the distribution of these parameters across different annotation categories. Proteins either got an annotation by MorF and/or a sequence based emapper search or not. (*“Annotation”* can entail a name or a description of the protein). After applying the Foldseek bit score cut-off, we gain a total of 16,285 proteins with annotations from MorF and emapper, 15,599 proteins without annotations, 8,914 proteins with MorF annotations only and 1,147 proteins with sequence based annotations only (Fig. S3A). Comparing different MorF parameters, it becomes clear again that large MSAs are helpful for gaining MorF based annotations (Suppl. Fig. S3B). Nevertheless, MSAs are not strictly necessary for getting MorF based annotations. Comparing pLDDTs it becomes obvious that proteins without any annotation usually have worse structural prediction quality (Fig. S3C) and vice versa.

Interestingly, proteins with only sequence based annotation (“No structure / Sequence”) have on average relatively high pLDDT values (Suppl. Fig. S3A). Nevertheless, their Foldseek bit score is too low (Suppl. Fig. S3D) to pass the bit score threshold for annotation. One possible explanation is that this category entails relatively short proteins (Suppl. Fig. S4A), which are shown to be suboptimal for structural searches [70], as they often do not pass (statistical) thresholds. On the other hand, proteins with MorF annotations have significantly higher Foldseek bit scores, reflecting the usefulness of the chosen bit score cut-off (Fig. S3D).

Comparing different annotation categories based on the distribution of physico-chemical parameters supports the statement above that they are not correlating with MorF parameters. Apart from molecular weight (Fig. S4B), none of the parameters such as gravy score, aromaticity index, isoelectric point, instability index and average flexibility score (Fig. S4B-F) differ significantly between the annotation categories.

**Fig S2.**
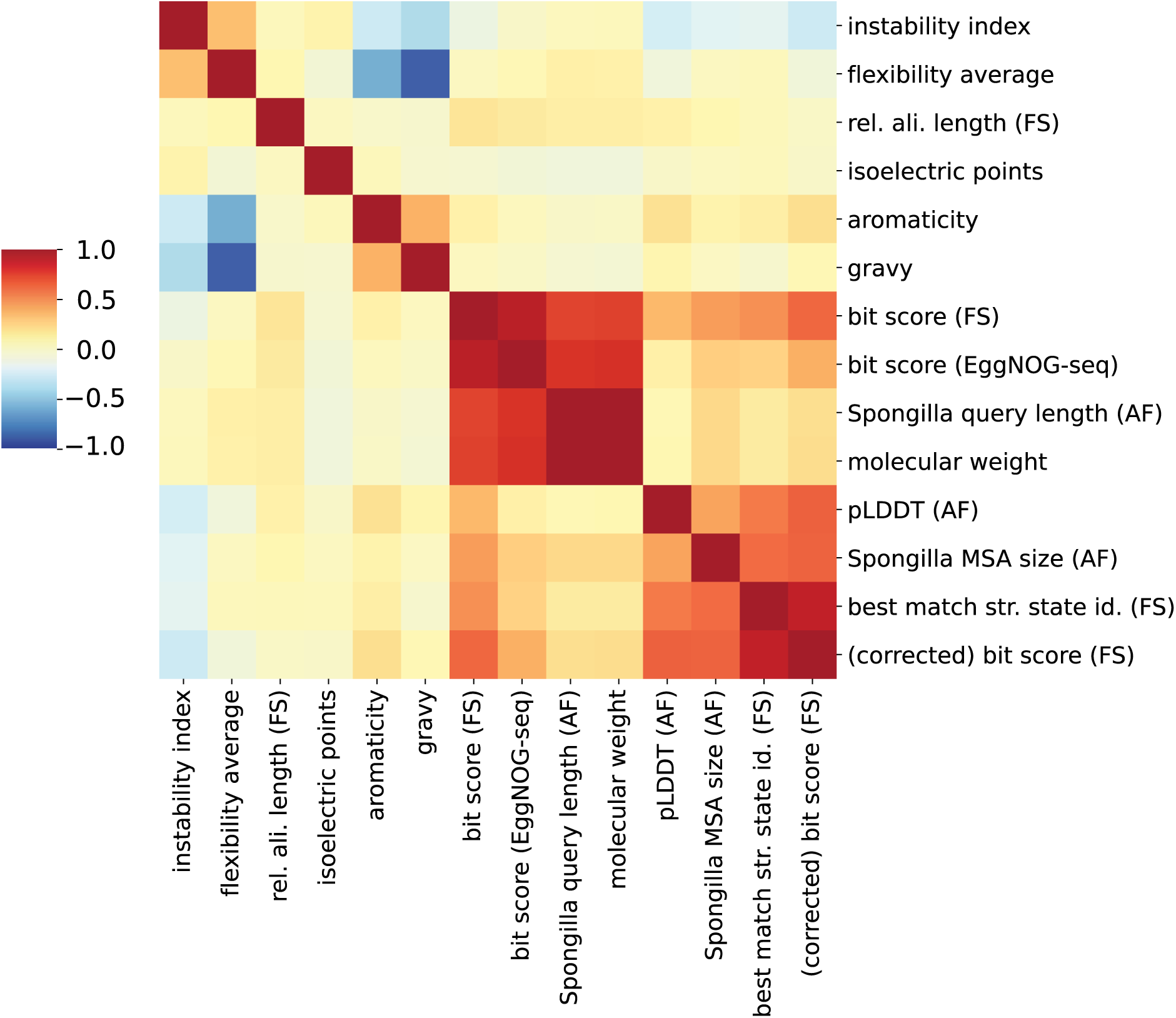
Pairwise correlation of metrics from the MorF pipeline as well as physico-chemical protein parameters. *instability index* : measure for proteins stability based on Guruprasad et al. [107]. *flexibility average* : average of structural flexibility scores across all amino acids of a protein based on Vihinen et al. [108]. *rel. ali. length (FS)*: percentage of the query (*Spongilla*) structure aligned with the best target structure. *isoelectric points* : prediction of the isoelectric point of a protein. *aromaticity* : measure for protein aromaticity calculated as the relative frequency of Phe/Trp/Tyr [109]. *gravy* : measure for the hydropathic caracter of a protein based on [110]. *bit score (FS)*: bit score of the Foldseek alignment between the query *Spongilla* structure and the structure of the best Foldseek target. *bit score (EggNOG-seq)*: bit score of the EggNOG alignment between the query *Spongilla* peptide sequence and the best target sequence. *Spongilla query length (AF)*: the query length of the input peptide. *molecular weight* : molecular weight of the protein. *plDDT (AF)*: average pLDDT of the predicted structure.

**Fig S3.**
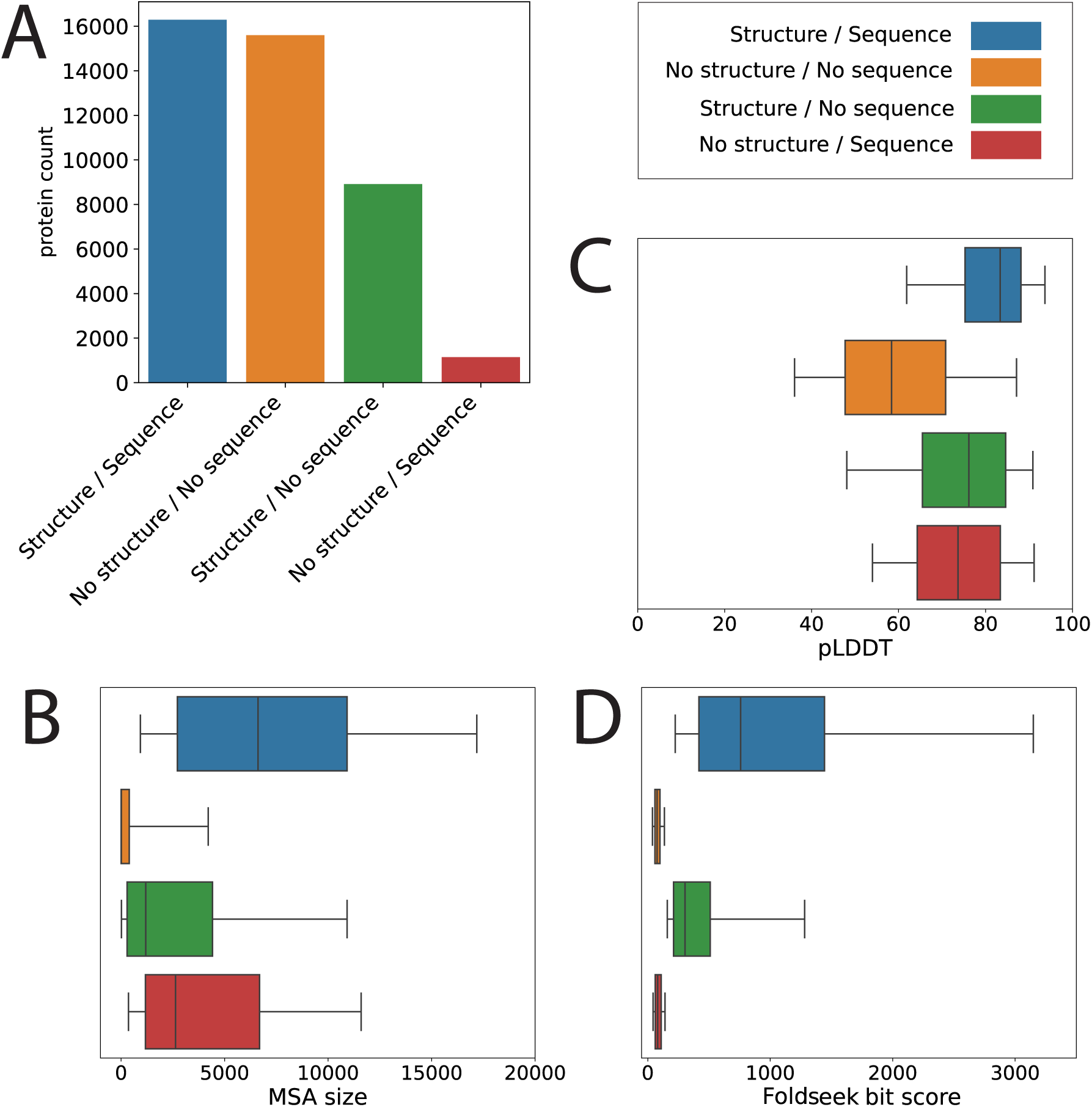
Comparison of MorF parameters in different annotation categories. A) Protein count in different annotation categories. Annotation includes either protein name or description. Protein counts are calculated *after* applying the Foldseek bit score cut-off. *“Structure/Sequence”* : Annotation by emapper sequence search as well as MorF *“No structure/No sequence”*: No annotation by emapper sequence search as well as MorF. *“Structure/No sequence”*: Annotation by MorF but no sequence based annotation by emapper. *“‘No structure/Sequence”*: Annotation by emapper sequence search but not by MorF. B) Distribution of AlphaFold MSA sizes across annotation categories. C) Distribution of AlphaFold pLDDTs across annotation categories. D) Distribution of Foldseek (uncorrected) bit scores across annotation categories.

**Fig S4.**
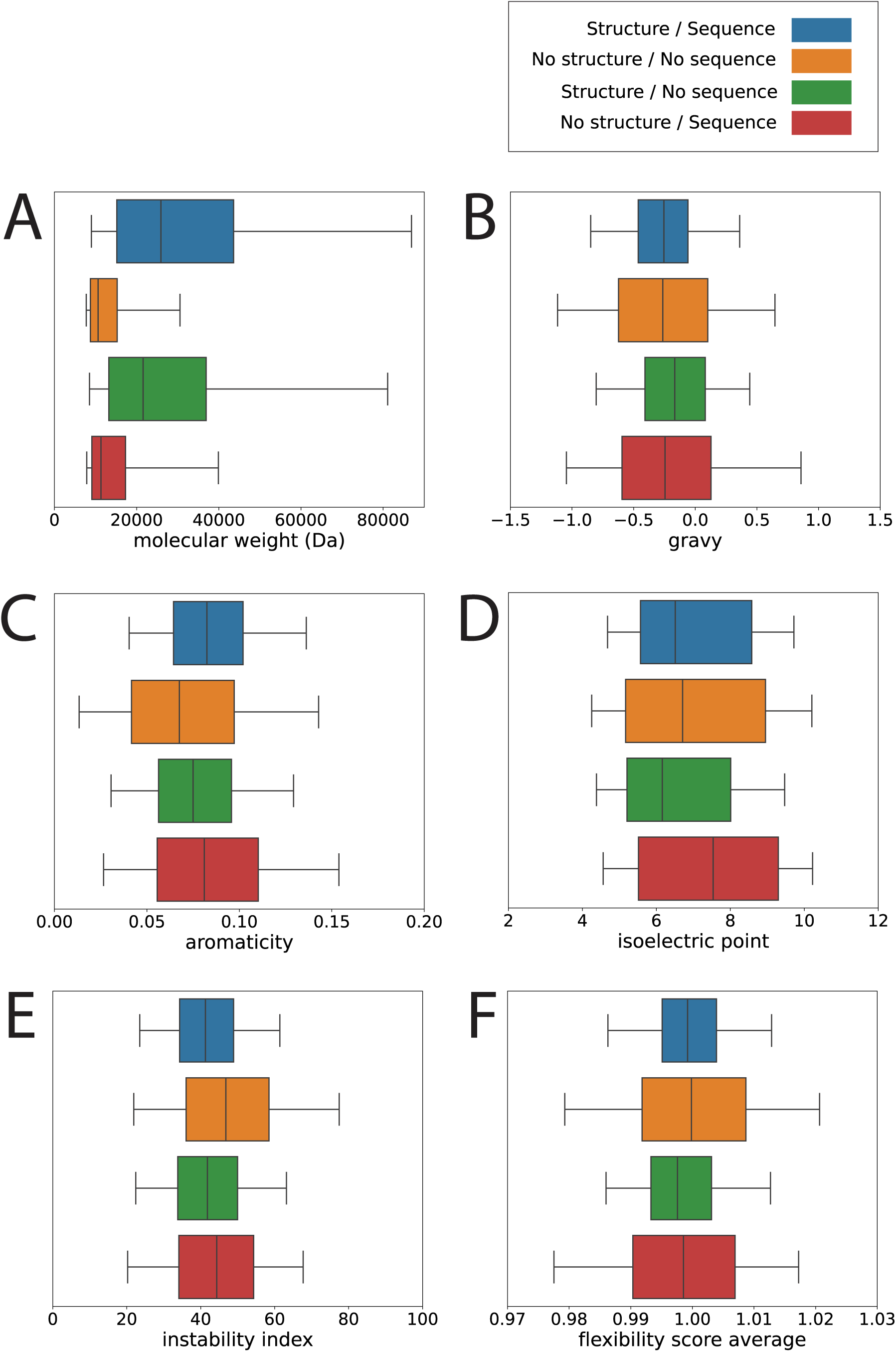
Comparison of physico-chemical protein parameters in different annotation categories. A) Distribution of molecular weights across annotation categories. B) Distribution of gravy scores across annotation categories. C) Distribution of aromaticity scores across annotation categories. D) Distribution of isoelectric points across annotation categories. E) Distribution of instability indices across annotation categories. F) Distribution of flexibility score averages across annotation categories.

*Spongilla MSA size (AF)*: size of the input MSA used for AlphaFold. *best match seq. id. (FS)*: structure state identity in the alignment between the query. *corrected bit score (FS)*: the Foldseek bit score divided by alignment length.

## C Comparison of MorF parameters between different agreement categories of structure and sequence based annotations

In order to explore and explain disagreements between structure and sequence based annotations, we compared different MorF parameters such as query length, MSA size, structure prediction quality as well as (alignment length) corrected Foldseek bit scores across different annotation agreement categories from (Fig. 1C; also refer to the corresponding notebook [16]). Most proteins do not have sequence (EggNOG) based annotations (Fig. S5A) and are excluded from this anaysis. This comparison shows that MSA sizes and corrected bit scores (Fig. S5E-F) are lower for proteins in the “no agreement” category compared to proteins with (partially) overlapping sequence and structure annotations. At the same time, the structure prediction quality for proteins (pLDDT) in the “no agreement” category are only marginally lower (Fig. S5C). Comparing Foldseek corrected bit scores across the annotation agreement categories broken down to multiple pLDDT buckets (Fig. S5B) shows that even in the cases of high prediction quality (*pLDDT* = 90 *−* 95%) the corrected Foldseek bit scores for the “no agreement” cases are very low, potentially leading to uncertainty in the structurally guided functional annotation. There are two potential reasons for this observation: First, Foldseek search results depend on the content of the structural databases that Foldseek searches against. Even very good structural predictions of sponge proteins can have low Foldseek bit scores if there are no morphologs present in the Foldseek database. With the ever expanding prediction of structures in the AlphaFold database, this issue will possibly be mitigated in the future. In fact, during the review process of this manuscript, a new version of AlphaFoldDB has been released, containing protein structures from all UniProt entries. Second, comparing query lengths of proteins in different annotation agreement levels (Fig. S5D), we observed a large proportion of long proteins in the “no agreement” category. Longer proteins often are composed of multiple domains that are linked via flexible linkers. Manual inspection showed that ColabFold correctly predicts the (globular) domains, leading to overall high pLDDT values, while predicting the flexible linker seemingly in a random fashion. This in turn leads to a random positioning of the domains relative to each other. This poses an issue for the subsequent Foldseek search in which only one of the domains can be superposed correctly, leading to the observed low corrected bit score. However, the extent to which this leads to incorrect functional annotation is still unknown and is outside the scope of the study.

Taking the possible explanations into account, sequence annotation would for now be preferable for proteins in the “no agreement” category. This agrees with our approach in the manuscript in which we primarily give preference to “legacy” sequence based annotation while supporting them with structural predictions and expand the annotation of those cases in which sequence is not sufficient for functional annotation.

**Fig S5.**
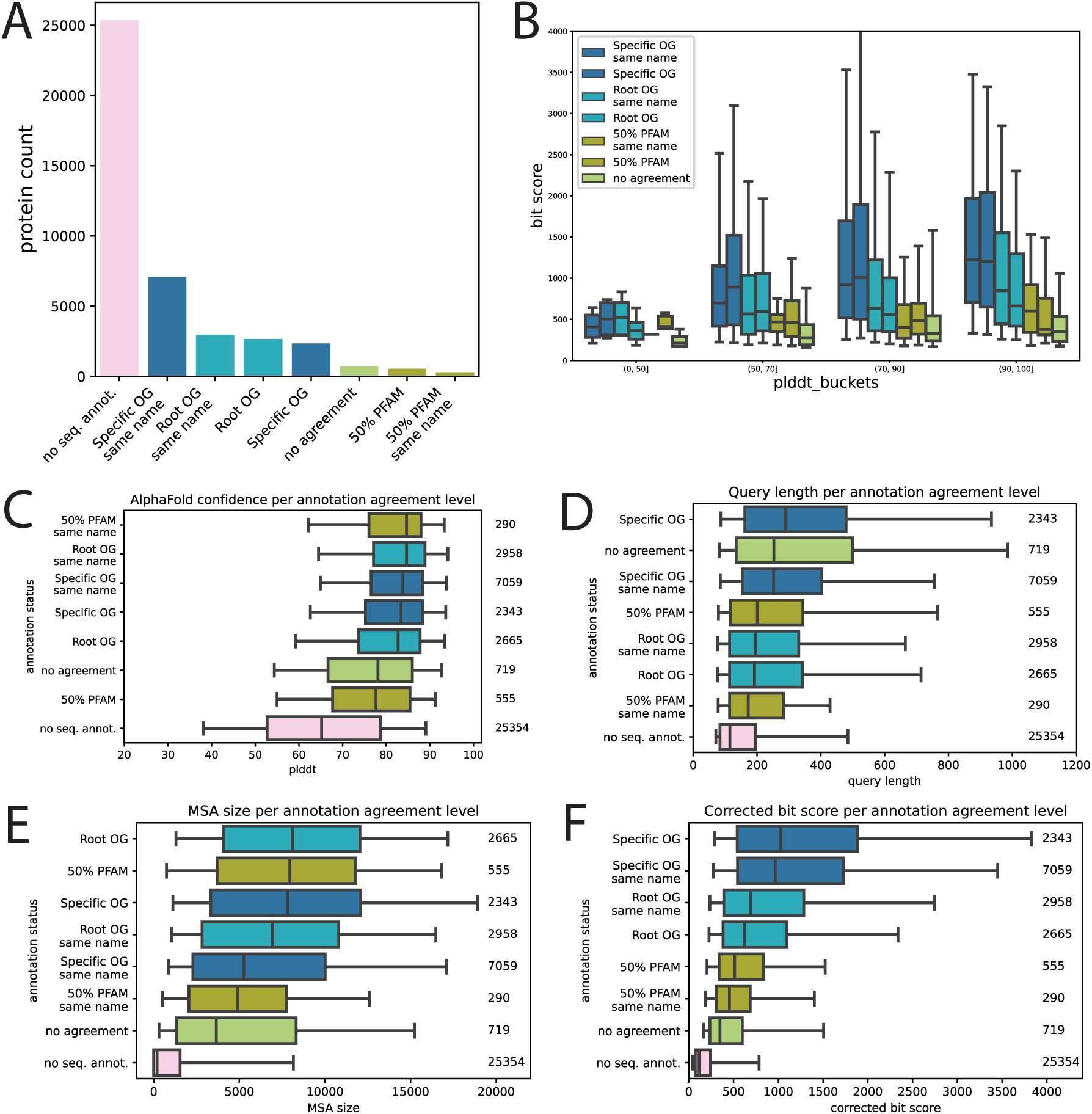
Comparison of MorF parameters between different categories of structure and sequence annotation agreements. A) Protein count of different annotation agreement categories. Most proteins (*∼*25, 000) do not have sequence (emapper) based annotations. 719 proteins do not agree in their sequence and MorF based annotations. B) Corrected (Foldseek) bit score comparisons across different annotation agreement categories within pLDDT value buckets. In all pLDDT buckets, proteins in the “no agreement” category show lowest average Foldseek bit scores. Structure prediction quality (pLDDT) (C), query length (D), MSA size (E) and corrected bit score (F) comparisons across annotation agreement categories.

## D Functional similarity of top morphologs

An indirect way to assess the performance of MorF at transferring functional annotations is to check how consistenly the top morphologs are assigned the same function. To do that, we define top morphologs via query-specific thresholds [111] and then compare the EC numbers of the top results to the best result [112], which in MorF would be assigned to the query protein.

We examined the AlphaFoldDB morphologs for each *Spongilla* query and calculated two thresholds to define the best morphologs. One is the multi-Otsu threshold [113], a technique from image analysis that tries to maximise between-class variance when thresholding pixel intensity values. The second threshold, referred to as “90th percentile”, was defined as the 90th percentile of the bit score range of each query’s putative morphologs. The 90th percentile threshold was much stricter than the multi-Otsu threshold in all cases.

We thresholded the AlphaFoldDB result matrix with the multi-Otsu cutoff for each query and queried UniProt via the Proteins API [114] to get the sequences of each protein in the filtered matrix. We submitted the sequences to the emapper webserver and removed hits with an e-value worse than 10^100^ to make sure only self-matches were kept. We populated the AlphaFoldDB result matrix with the EggNOG annotation and removed entries that didn’t have an EC number. We then additionally removed all queries that only had one possible target remaining.

We then assessed the functional overlap of the Otsu-thresholded and 90th percentile-thresholded morphologs with the top morpholog. We quantified this as the average agreement of the EC numbers of the morphologs with the EC number of the top morpholog. An average agreement of 3 would mean that the top morphologs, on average, share the first three positions of the EC number of the top morpholog.

A total of 24,380 queries had at least two morphologs with an EC clearing the multi-Otsu threshold, with an average of 13.74 top morphologs per query. A total of 7,072 queries had at least two morphologs with an EC clearing the much stricter 90th percentile threshold, with an average of 7.37 top morphologs per query.

The Otsu threshold seemed to be too permissive, with a large majority of the queries showing significant divergence within their top morphologs (average EC overlap of 2.02, so barely within the same enzyme subclass). On the other hand, in the 90th percentile threshold top morphologs had an average EC overlap of 3.7 - close to complete agreement. In the same subset of proteins (90th percentile threshold), the first and second best morphologs have a different EC in 10.88% of cases.

For the narrow subset of proteins we could feasibly examine, we conclude that the top morphologs of each protein overwhelmingly have the same function, offering another indication that MorF’s approach is fundamentally valid.

## E GO term annotation comparison

**Fig S6.**
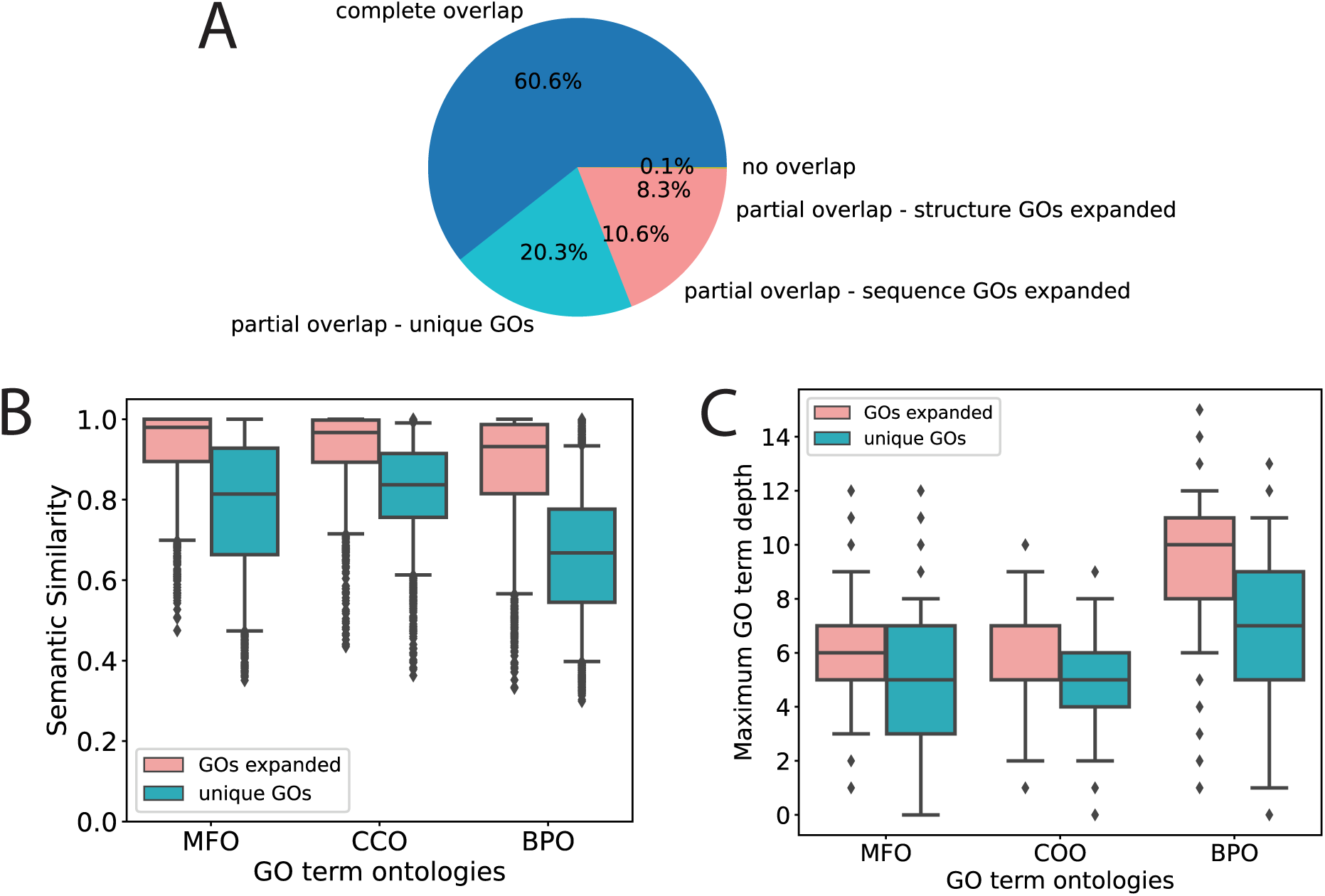
Comparison of GO term assignment, semantic similarity and depth between structure- and sequence based annotations. A) Overlap of GO term assignments between MorF and sequence based annotations. (“complete overlap”: GO term assignments are identical; “partial overlap - unique GOs”: GO terms overlap with unique GO terms assigned to both, MorF and sequence based annotations; “partial overlap - sequence GOs expanded”: additional GO terms assigned by sequence based annotation; “partial overlap - structure GOs expanded”: additional GO terms assigned by MorF; “no overlap”: no overlap between GO term assignments (17 cases total)) B) GO term semantic similarities between GO term assignments of the “partial overlap” categories across GO term ontologies (“MFO”: molecular function, “CCO”: cellular component, “BPO”: biological process) C) Maximum depths of *overlapping* GO terms between sequence- and structure based annotations across GO term ontologies. Interpretation of GO term depths is challenging as different depths might imply different levels of details for different branches in the hierarchy.

**Table 2.** High confidence structures with non-helical appearance. The last column contains a summary of the significant BLASTp results in the nr database. Question marks denote peptides who did not find any named sequence homologs or significant PFAM domain hits.

| isoform | q len | #MSA | %seq. id. | bit score | plddt | BLASTp |
| --- | --- | --- | --- | --- | --- | --- |
| c88419.g1.i1 | 154 | 462 | 0.337 | 580.0 | 85.01 | CRN5 |
| c104256.g1.i1 | 375 | 163 | 0.272 | 785.0 | 82.42 | CRN5 |
| c104256.g1.i2 | 299 | 184 | 0.267 | 781.0 | 90.35 | CRN5 |
| c104256.g1.i3 | 336 | 167 | 0.262 | 777.0 | 88.39 | CRN5 |
| c104256.g1.i4 | 332 | 158 | 0.263 | 796.0 | 90.26 | CRN5 |
| c104256.g1.i6 | 253 | 196 | 0.275 | 774.0 | 92.81 | CRN5 |
| c94352.g2.i1 | 127 | 1,725 | 0.500 | 299.0 | 86.06 | DD3-3 |
| c89711.g1.i1 | 144 | 2,424 | 0.441 | 380.0 | 86.73 | DD3-3 |
| c89711.g2.i1 | 75 | 2,340 | 0.611 | 210.0 | 77.48 | DD3-3 |
| c112476.g1.i1 | 179 | 1,450 | 0.352 | 434.0 | 83.19 | DD3-3 |
| c103626.g1.i1 | 670 | 3,392 | 0.324 | 2224.0 | 91.33 | DD3-3 |
| c103626.g1.i2 | 670 | 3,424 | 0.329 | 2258.0 | 91.18 | DD3-3 |
| c103626.g2.i1 | 667 | 3,427 | 0.326 | 2215.0 | 90.01 | DD3-3 |
| c102584.g1.i1 | 578 | 3,328 | 0.355 | 2091.0 | 81.55 | DD3-3 |
| c102584.g1.i2 | 187 | 1,994 | 0.416 | 522.0 | 89.04 | DD3-3 |
| c101972.g1.i1 | 778 | 3,481 | 0.378 | 2536.0 | 85.58 | DD3-3 |
| c101972.g1.i2 | 778 | 3,481 | 0.378 | 2534.0 | 85.58 | DD3-3 |
| c101972.g1.i3 | 778 | 3,481 | 0.378 | 2537.0 | 85.54 | DD3-3 |
| c104453.g2.i1 | 359 | 368 | 0.177 | 662.0 | 75.80 | fol. epith. yolk prot. subunit |
| c104453.g2.i2 | 280 | 356 | 0.172 | 658.0 | 89.90 | fol. epith. yolk prot. subunit |
| c104453.g2.i3 | 340 | 378 | 0.185 | 634.0 | 78.61 | fol. epith. yolk prot. subunit |
| c104453.g2.i4 | 236 | 374 | 0.183 | 594.0 | 83.73 | fol. epith. yolk prot. subunit |
| c104453.g3.i1 | 182 | 341 | 0.117 | 298.0 | 70.54 | fol. epith. yolk prot. subunit |
| c98985.g1.i2 | 73 | 1,852 | 0.138 | 45.0 | 70.12 | XPO5 (Exportin-5) |
| c88005.g3.i1 | 130 | 3,213 | 0.274 | 363.0 | 94.66 | hemopexin repeat-containing protein |
| c70965.g1.i1 | 169 | 299 | 0.209 | 354.0 | 77.62 | pinocchio |
| c103612.g1.i1 | 89 | 1,253 | 0.152 | 46.0 | 73.88 | kinesin-like protein KIF23 |
| c102197.g1.i3 | 312 | 257 | 0.179 | 649.0 | 86.28 | bamB |
| c100331.g2.i1 | 281 | 43,509 | 0.219 | 821.0 | 82.62 | hydralysin-2-like |
| c93555.g2.i2 | 77 | 439 | 0.109 | 80.0 | 87.77 | ? |
| c86044.g1.i1 | 87 | 7,052 | 0.077 | 90.0 | 88.87 | ? |
| c83472.g1.i1 | 98 | 553 | 0.177 | 90.0 | 84.77 | ? |
| c108675.g1.i1 | 73 | 3,174 | 0.224 | 51.0 | 78.17 | ? |
| c100701.g1.i2 | 69 | 1 | 0.123 | 61.0 | 81.77 | ? |
| c100429.g1.i1 | 79 | 43,996 | 0.152 | 85.0 | 71.74 | ? |

## F Differentially expressed genes per cell type

**Fig S7.**
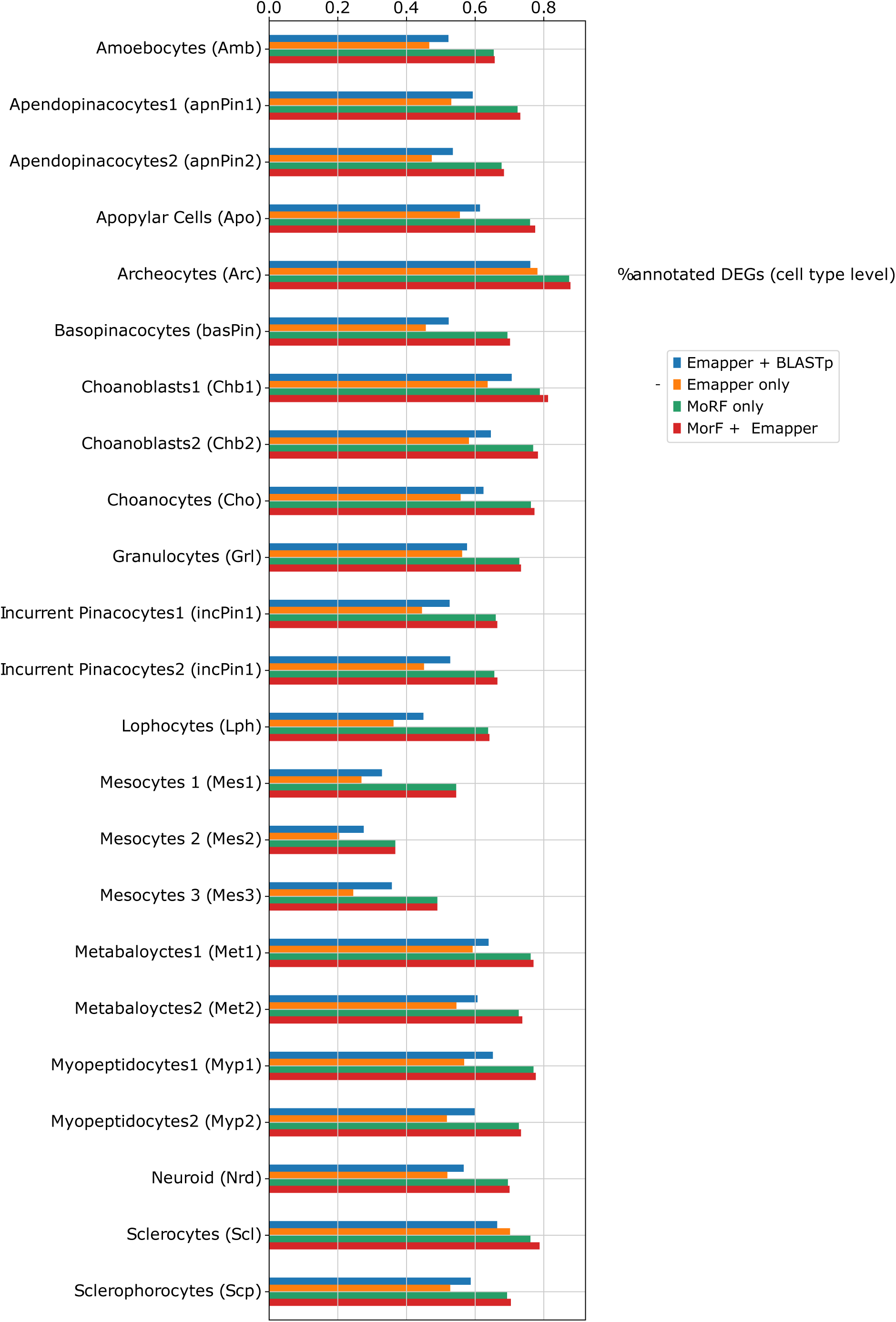
The percentage of marker genes per cluster as per Musser *et al.* that are annotated by different methods.

## G Mesocyte marker gene expression

**Fig S8.**
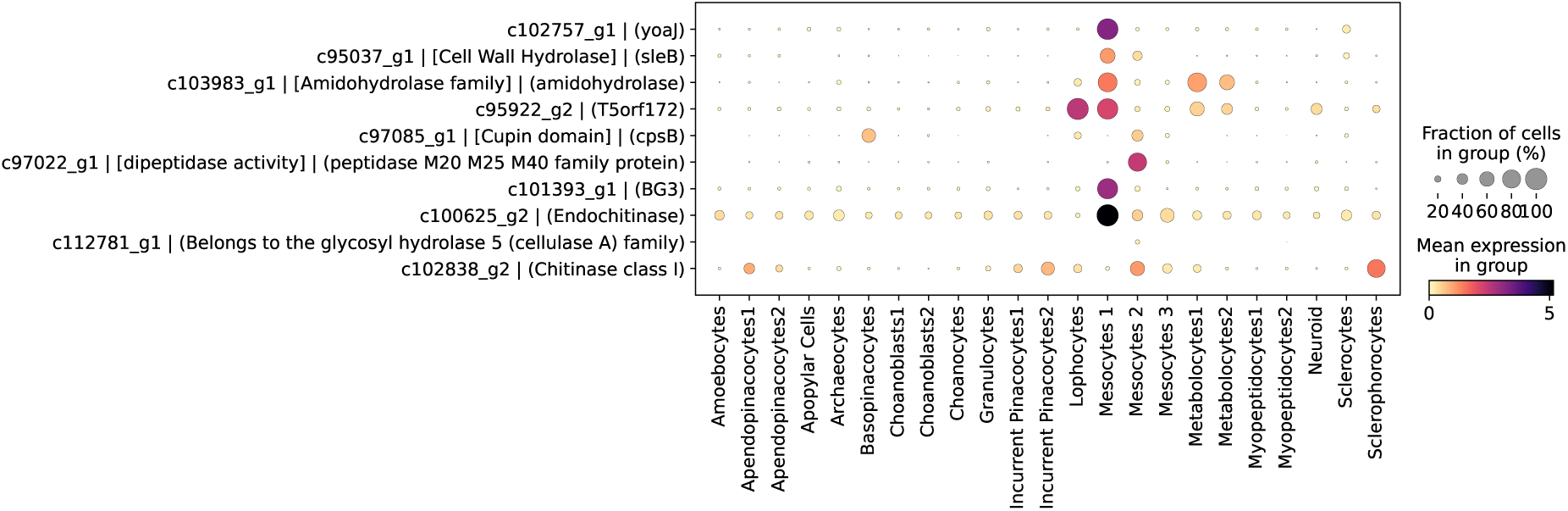
Mesocyte marker gene expression. Dotplot of mesocyte marker gene expression for HGT candidates. emapper annotation in square brackets; MorF annotation in round brackets. Mean expression of log-transformed, normalized counts.

## H Structure-sequence agreement in model species

As proof of principle we explored the power of structural similarity to identify orthologous genes for a number of model species whose proteomes already had predicted structures in AlphaFoldDB. We aligned AlphaFoldDB against itself using Foldseek. For each query protein we kept the best target from outside the taxonomic group and compared the eukaryotic orthogroup assignment. The best structural hit overwhelmingly belonged to the same orthogroup as the query (Suppl. Table 3).

We used the Proteins API [114] to obtain the species name for each entry [115]. We decorated each entry with its orthogroup assignment from the EggNOG database [18] (v5.0). Not all proteins currently in AlphaFoldDB could be assigned an orthogroup; we ignored comparisons where the query or its best target were missing the orthogroup assignment. The full table can be found in the corresponding notebook [116].

We compared the remaining eligible cases and found overwhelming agreement between the eukaryotic orthogroup of the best structural target and the orthogroup of the query protein, confirming that structural similarity can, in principle, detect homology relationships. Though the available targets were restricted to be outside the clade, order, phylum, or even kingdom, top morphologs still are homologs in the large majority of cases, a very encouraging indication that MorF is able to detect homology over long evolutionary distances.

As a more demanding application, we compared enzyme function between distantly related species to see how often function was correctly predicted by structural similarity (see corr. notebook [117]). In particular, we were interested in cases where homology via sequence similarity was no longer detectable but the EC number still overlapped.

We performed two comparisons between well-studied, distantly related eukaryote species: baker’s yeast against human, and *Arabidopsis thaliana* against human. We downloaded the predicted protein structures from the AlphaFold protein database, performed structural alignment against AlphaFoldDB, and then filtered out non-human hits. Using the EggNOG annotation of the query and target proteins, we removed pairs that belonged to the same protein families (same root orthogroup or same most specific orthogroup; opisthokont for human/yeast and eukaryote for *A. thaliana*.)

The best morphologs of yeast enzymes without previously detected homologs in human agree on all four positions of the EC number in 53/146 (36%) cases. Broader similarity (three of four EC positions) can be found in 109/146 (75%) cases. Similarly, the best morphologs of *A. thaliana* enzymes without clear homologs in human have the same EC in 176/532 (33%) cases, and share the first three digits in 357/532 (67%) cases.

**Table 3.** Structural similarity correctly identifies homologs for model species in the absence of close relatives. Columns: total number of queries with a hit outside the species context; total number of cases where both query and best non-species target are annotated by EggNOG; percentage of these cases where the eukaryote orthogroup between query and target is the same; taxonomic group. Species that belong to the same taxonomic group were excluded when performing this comparison.

| species | #queries with<br>out-of-group<br>targets | #eligible<br>queries | %eligible<br>agreement | taxonomic<br>group |
| --- | --- | --- | --- | --- |
| *H. sapiens* | 20,392 | 13,838 | 74.30% | Vertebrata |
| *M. musculus* | 21,507 | 15,084 | 75.05% | Vertebrata |
| *R. norvegicus* | 19,246 | 13,180 | 74.60% | Vertebrata |
| *D. rerio* | 24,626 | 13,303 | 75.36% | Vertebrata |
| *D. melanogaster* | 13,418 | 8,828 | 79.69% | Arthropoda |
| *C. elegans* | 19,599 | 11,177 | 67.29% | Nematoda |
| *S. mansoni* | 13,845 | 1,730 | 91.85% | Nematoda |
| *A. thaliana* | 27,357 | 19,800 | 83.78% | Eudicots |
| *G. max* | 55,759 | 32,239 | 85.40% | Eudicots |
| *Z. mays* | 38,818 | 18,644 | 85.47% | Monocots |
| *O. sativa* subsp. *japonica* | 41,794 | 20,306 | 86.51% | Monocots |
| *S. cerevisiae*<br>(str. ATCC 204508/S288c) | 6,013 | 4,347 | 83.41% | Saccharomycetales |
| *S. pombe* (str. 972/<br>ATCC 24843) | 5,104 | 3,982 | 90.51% | Saccharomycetales |
| *C. albicans* (str. SC5314/<br>ATCC MYA-2876) | 5,973 | 4,118 | 84.41% | Saccharomycetales |
| *C. carrionii* | 11,169 | 7,040 | 81.29% | Chaetothyriales |
| *P. lutzii* (str. ATCC MYA-826/Pb01) | 8,791 | 5,086 | 84.19% | Eurotiomycetes |
| *S. schenckii* (str. ATCC 58,251/<br>de Perez 2211183) | 8,652 | 6,349 | 78.41% | Sordariomycetes |
| *T. brucei brucei*<br>(str. 927/4 GUTat10.1) | 8,476 | 4,303 | 60.26% | Trypanosomatida |
| *T. cruzi* (str. CL Brener) | 19,026 | 8,782 | 56.56% | Trypanosomatida |
| *L. infantum* | 7,914 | 4,423 | 56.14% | Trypanosomatida |
| *D. discoideum* | 12,584 | 6,312 | 63.96% | Amoebozoa |

**Fig S9.**
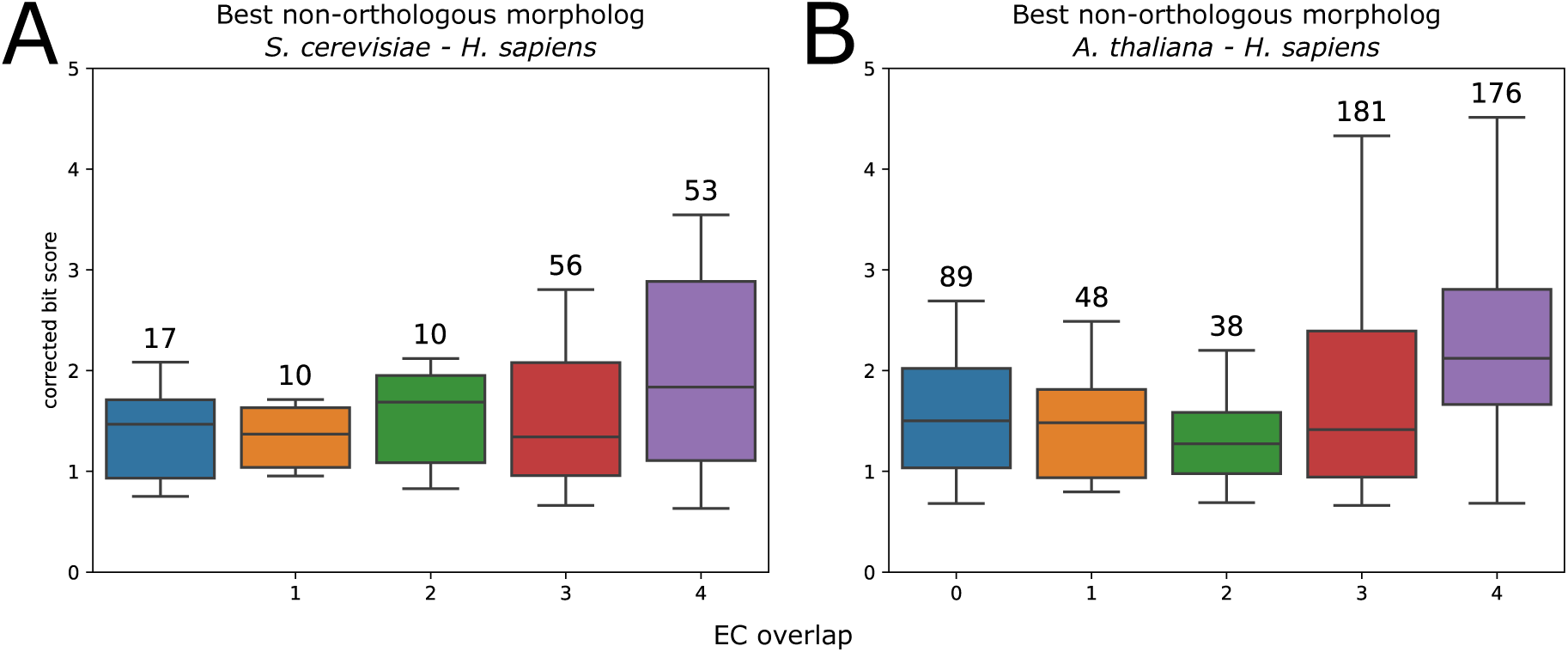
Remote species comparison: Distribution of the EC overlap of the best non-orthologous morphologs between distantly related species, against corrected bit score. For the majority of cases, same structure means a significant overlap in EC despite the absence of orthology. While high EC overlap usually means a higher corrected bit score (bit score divided by alignment length), it is not always the case.

## I Comparison to sequence-profile searches

To gain a better idea of the limits of structural similarity as a means to transfer functional annotation we compared MorF’s performance to that of HMMER [37]. In particular, we used eggnog-mapper in “HMMER” mode and performed sequence-to-profile searches for the translated *S. lacustris* proteome against the eukaryotic HMMs of the EggNOG database.

Emapper-hmmer finds putative homologs for 28,897 proteins, compared to 17,990 by standard emapper and 25,232 morphologs found by by MorF (Fig. S10B, lower left).

The additional sensitivity of emapper-hmmer comes at the cost of precision, an expected trade-off, especially when using sequence profiles at the eukaryote level. Concretely, this means that we expect emapper-hmmer to return orthogroups from higher taxonomic levels more often than emapper or MorF, and therefore, in average, less orthogroups. It also means that we expect emapper-hmmer to fail to assign a name or a description to many annotations, as the taxonomic unit where orthology is detected will be more vague than, e.g. a gene family. Since protein names, despite their drawbacks, are the most common human-readable short descriptors of function, this is a major drawback for the use of sequence profiles for functional annotation at scale, such as the use case we present here.

Following our expectations, MorF results have significantly more orthogroups per query than either emapper (two-sample Kolmogorov-Smirnov test; statistic 0.56, p-value 0) or emapper-hmmer (two-sample Kolmogorov-Smirnov test; statistic 0.57, p-value 0) . Surprisingly, emapper-hmmer results have slightly more orthogroups per query than emapper, though with low statistical support (two-sample Kolmogorov-Smirnov test; statistic 0.057, p-value 6.9*^−^*^24^) (Fig. S10A). In the same vein, 11,700 of the emapper-hmmer annotations don’t have a preferred name, compared to 5,200-5,300 for MorF/standard emapper. Similarly, emapper-hmmer doesn’t even produce a description for ca. 2,600 proteins, compared to 38 for MorF and ca. 600 for standard emapper.

**Fig S10.**
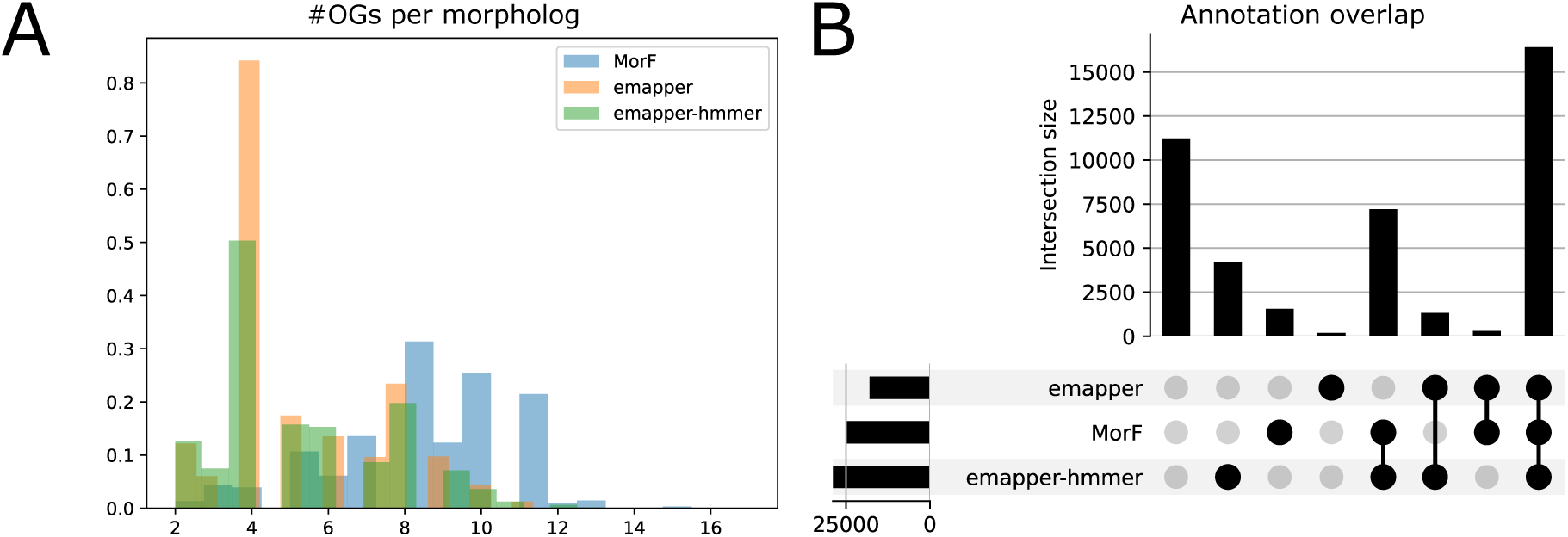
Comparison of MorF to sequence-based annotation. A) Histogram of the number of EggNOG orthogroups assigned to each annotated query. As orthogroups have hierarchical relationships, more orthogroups generally indicate a more precise annotation. Mean and standard deviation: emapper (5.23, 2.08), emapper-hmmer (5.35, 2.23), MorF (8.28, 5.21). B) Breakdown of overlaps between different annotation strategies. The upper panel shows the number of *Spongilla* proteins annotated by emapper, MorF, emapper-hmmer, or their combinations. The lower right panel shows the categories being combined, and the lower left panel shows the total number of annotations produced by each tool.

At the same time, we examined the agreement between emapper-hmmer and MorF, similar to the comparison of MorF to emapper at the task of homology detection. For compatibility purposes, we focused on the 16,346 proteins that were annotated by all three methods (Fig. S10B). Since emapper-hmmer was run on the eukaryotic orthogroup profiles we added a comparison at the eukaryote orthogroup level. In the vast majority of cases MorF produces the same homology assignment as emapper-hmmer.

Furthermore we repeated the analysis presented in Fig. 1D, now including emapper-hmmer (Fig. S11). Reflecting the results that were already mentioned, emapper-hmmer performs better than standard emapper. However, when it comes to the marker genes that will drive biological discovery, MorF still outperforms sequence similarity as an annotation tool.

**Table 4.** Percentage of proteins where annotation agreed between the tools at different taxonomic levels. Where not explicitly noted, all 16,346 proteins were eligible for comparison.

| tool combination | most specific OG | eukaryotic OG (eligible) | root OG |
| --- | --- | --- | --- |
| emapper / emapper-hmmer | 66.74% | 86.25% (16,032) | 91.15% |
| MorF / emapper-hmmer | 48.69% | 82.06% (15,889) | 87.18% |
| MorF / emapper | 56.77% | 89.11% (15,745) | 90.71% |

**Fig S11.**
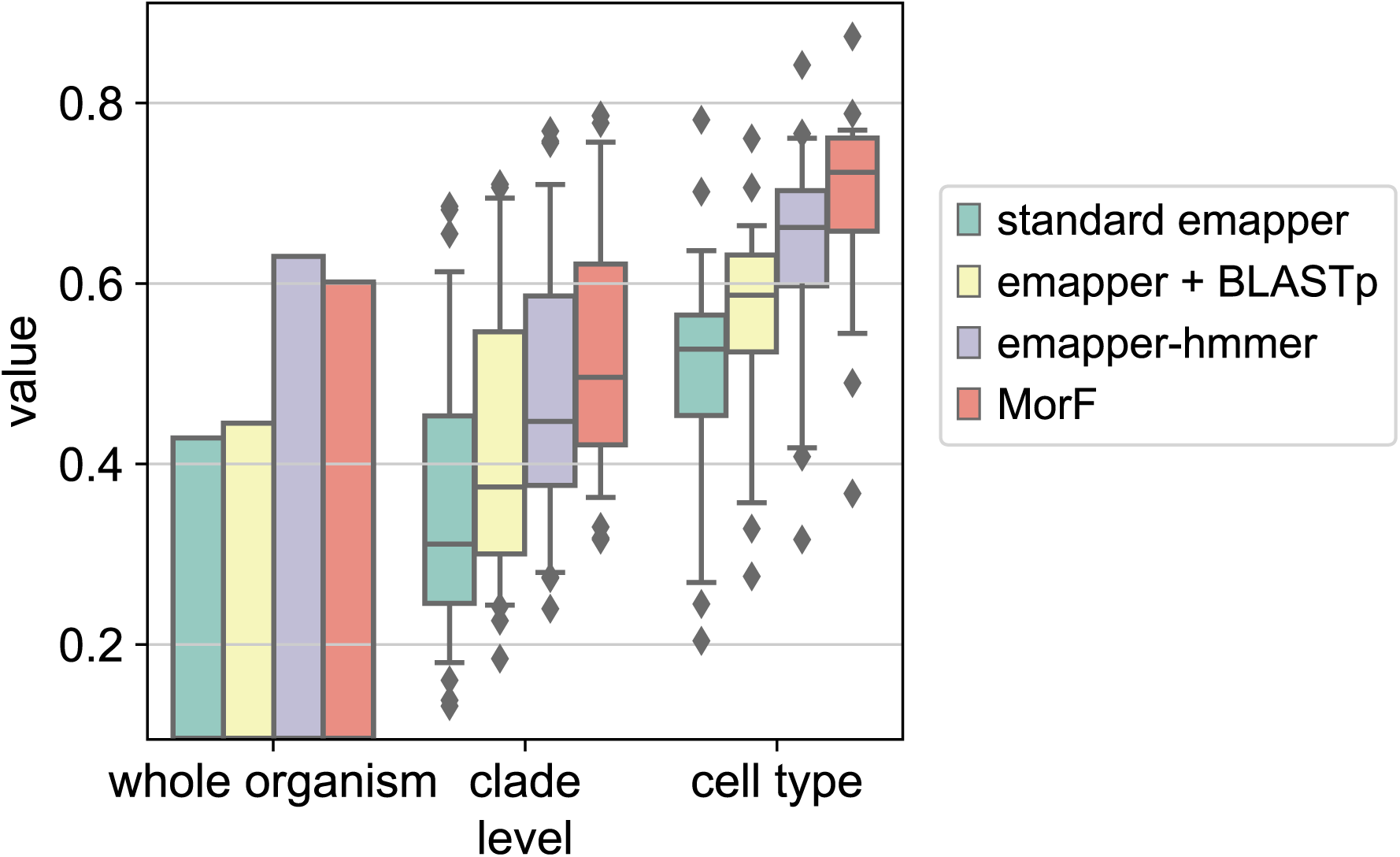
Proportion of annotated proteins for various relevant groups. “Whole organism” denotes the entire proteome; “clade level” denotes the marker genes for cell type families in the single-cell data of Musser *et al.* [9], while “cell type” denotes the marker genes for the various cell types in the same dataset.

In conclusion, for the task of producing usable functional annotation, MorF compares favorably to emapper-hmmer, a state-of-the-art tool for profile-sequence searches. It proves capable of largely reproducing the same orthology assignments, and annotates a higher proportion of cell type-specific genes, thereby aiding biological discovery.

**Table 5.**
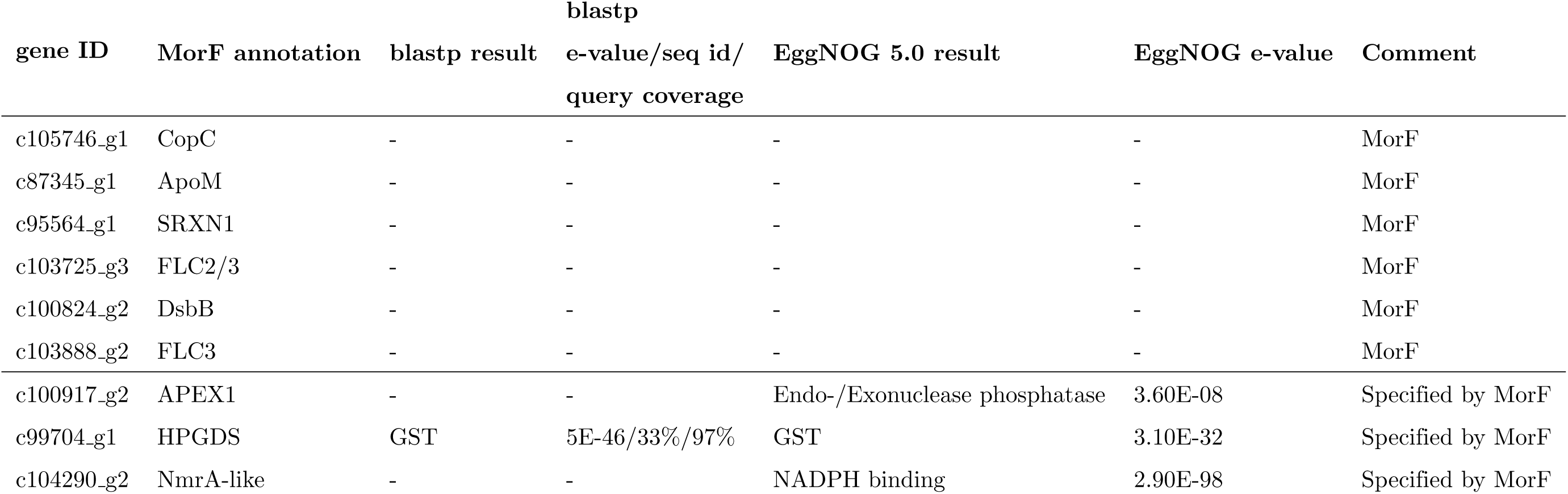
Blastp and EggNOG search results of newly identified myopeptidocyte marker gene morphologs. Blastp searches were run against non redundant protein database (nr) in default mode. EggNOG sequence searches were run in default mode. E-value, sequence identity (seq id) and query coverage are reported from the best blastp hit or EggNOG orthology group.

## J Myopeptidocyte marker genes

**Table 6.**
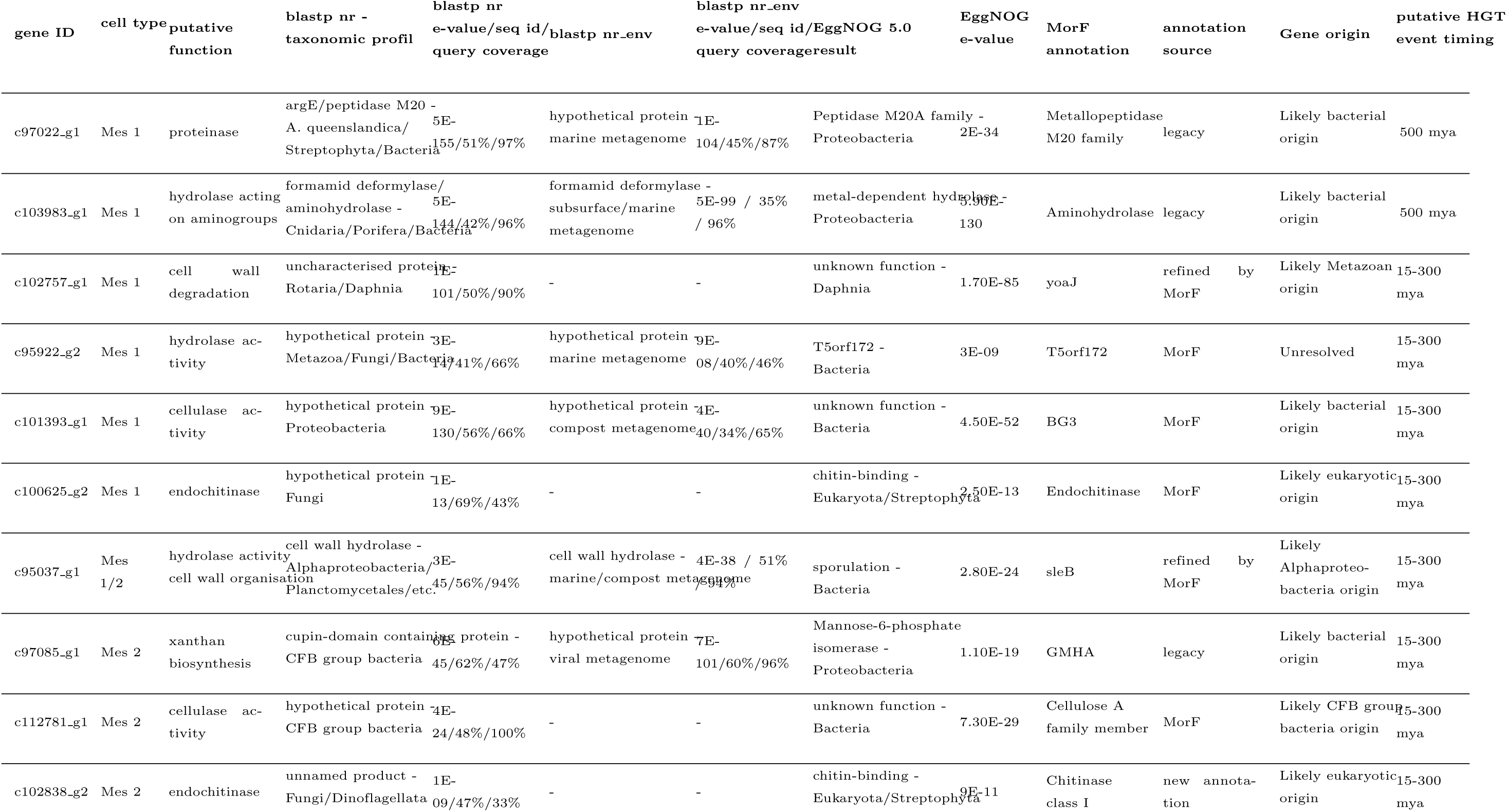
Blastp and EggNOG search results of newly identified mesocyte marker gene morphologs. Blastp searches were run against non redundant protein database (nr) as well as environmental metagenome databases (env nr) in default mode. EggNOG sequence searches were run in default mode. E-value, sequence identity (seq id) and query coverage are reported from the best blastp hits or best scoring EggNOG orthologous group. Based on the blastp and EggNOG results we hypothesised plausible gene origins. Potential timing of putative HGT events are based on the distribution of the genes within Demosponge phylogeny [118].

## K Mesocyte marker genes

## L Presence of sponge HGT candidates in Choanoflagellates

To gain a more complete picture about the putative HGT events suggested by MorF we performed sensitive sequence searches (MMseqs2, setting -s 7.0; see [119]) of the *Spongilla* HGT candidates against the refernce proteomes of emerging Choanoflagellate models *S. rosetta* and *M. brevicollis*. We find *Spongilla* genes c103983 g1 and c97022 g1 in both species, with rather low e-values. This is intriguing, as all the *S. rosetta* targets had already been identified as putative HGT results in earlier work [63]; in conjunction with their broad phylogenetic distribution in sponges it is tempting to speculate that these are ancient HGT events that occurred before the split of Choanoflagellates and sponges.

